# Artefact or ecology? Integrating bacteria-phage defence and mobilome signatures to distinguish human-associated MAGs from probable contaminants in low biomass cancer metagenomes

**DOI:** 10.64898/2026.11.27.769564

**Authors:** Vigneshwaran Muthuraman, Vikki Rand, Saadlee Shehreen

**Author notes:** **Corresponding author and email** Saadlee Shehreen.

## Abstract

Low-biomass cancer metagenomes are highly susceptible to contamination, complicating the identification of authentic human-associated microorganisms. Here, we investigated whether bacteria– phage co-evolutionary signatures and mobile genetic elements provide complementary evidence for distinguishing human-associated bacteria from probable contaminants in cancer metagenomes. We analysed 514 metagenome-assembled genomes (MAGs) from metastatic tumour biopsies and 785 MAGs from an independent colorectal cancer stool cohort, integrating phage–host predictions, defence and anti-defence systems, insertion sequences, plasmids, prophages and genome architecture. Of the 514 tumour-derived MAGs, 131 (25.5%) were classified as probable contaminants based on concordant taxonomic assignments and ecological signatures indicative of non-human-associated origins. Human-associated MAGs exhibited concordant phage–host affiliations, conserved defence repertoires and distinct mobilome profiles across cohorts. Probable contaminants displayed larger genomes, higher GC content, greater insertion-sequence abundance and approximately 20-fold higher plasmid-associated sequence burdens. Although most shared defence signatures reflected bacterial lineage, AbiQ and Wadjet-III emerged as notable exceptions, being detected in both cancer-derived datasets while remaining uncommon in reference genomes and absent from probable contaminants. Together, these findings support the use of complementary phage–host, defence and mobilome signatures to assess the ecological plausibility of microbial genomes recovered from cancer metagenomes. They also identify candidate defence systems for experimental investigation, with potential relevance to future defence-guided phage selection in microbial modulation.

**Repositories:**

1. Wirbel colorectal cancer metagenome-assembled genomes (1) generated in this study (785 MAGs): Zenodo, https://doi.org/10.5281/zenodo.24062335
2. Analysis code and processed data: GitHub, https://github.com/vikos77/Cancer-Microbiome, archived at Zenodo, https://doi.org/10.5281/zenodo.24060765
3. Hartwig MAGs (2): Zenodo, https://doi.org/10.5281/zenodo.10777510
4. Wirbel CRC raw metagenomic reads: European Nucleotide Archive, PRJEB6070 and PRJEB27928

**Impact statement:** Low-biomass tumour metagenomes are highly prone to exogenous contamination, complicating the verification of authentic host-associated bacteria. Integrating phage–host signatures, anti-phage defence repertoires, and mobilome profiles across cancer metagenome-assembled genomes can reliably distinguish genuine human-associated taxa from contaminants.

**Data summary:** The authors confirm all supporting data, code and protocols have been provided within this article or through supplementary data files.

1. Hartwig MAGs. The 514 metagenome-assembled genomes (MAGs) analysed here were assembled and binned from Hartwig Medical Foundation metastatic tumour biopsies (https://doi.org/10.1016/j.cell.2024.03.021) (2). MAG sequences were obtained from that study’s Zenodo deposit (https://doi.org/10.5281/zenodo.10777510) and MAG quality metrics from its Table S2.
2. Wirbel colorectal cancer (CRC) cohort. Raw paired-end shotgun metagenomic reads for 60 CRC stool samples were downloaded from the European Nucleotide Archive (ENA) under study accessions PRJEB6070 (38 samples, https://www.ebi.ac.uk/ena/browser/view/PRJEB6070) and PRJEB27928 (22 samples, https://www.ebi.ac.uk/ena/browser/view/PRJEB27928). The 785 MAGs assembled from these reads, with a per-MAG manifest giving source sample, contig count, genome length and MD5 checksum, are available at Zenodo (https://doi.org/10.5281/zenodo.24062335).
3. Code and processed data. The analysis code and the compiled annotation-tool outputs are available at https://github.com/vikos77/Cancer-Microbiome and archived at Zenodo (https://doi.org/10.5281/zenodo.24060765). This release also lists the accessions and assigned isolation source of 3,248 complete RefSeq genomes used for comparison (data/reference_genomes/defensefinder_summary_refgenome.tsv).
4. Supplementary material. Tables S1-S6 are provided as one Excel file, and the Supplementary Methods and Figures S1 and S2 as one PDF.

## 5. Introduction

Cancer-associated microbiomes have attracted growing interest because of their potential roles in tumour biology, immune responses and treatment outcomes. Reported associations between microbial composition, tumour hypoxia and responses to immunotherapy suggest that microorganisms may contribute to clinically relevant differences between patients(2). However, the origin and biological significance of microbial signals recovered from cancer metagenome datasets remain debated(3, 4). A central challenge is establishing whether these signals represent authentic human-associated microorganisms, transient microbial material or technical artefacts(2).

This challenge is particularly acute in low-biomass tumour samples, where small amounts of contaminating DNA can substantially influence the apparent microbial composition. Contamination introduced during sample collection, DNA extraction or sequencing, together with incomplete removal of human sequences and errors in taxonomic assignment, can generate misleading microbial profiles(5–7). Taxonomic identification alone is therefore insufficient to distinguish human-associated bacteria from probable contaminants. Negative controls, rigorous computational filtering and independent validation remain essential, but complementary evidence grounded in microbial ecology such as phage-host interactions could strengthen the interpretation of these datasets.

Repeated encounters between bacteria and bacteriophages leave genomic traces, including integrated prophages, phage-derived spacers within CRISPR arrays and mechanisms of resistance to infection(8–10). These features record aspects of microbial interaction histories that are not captured by taxonomic profiles alone. Phages also contribute to horizontal gene transfer (HGT), alongside other mobile genetic elements (MGEs), including plasmids, transposons and integrative elements. By mediating the exchange of genetic material, MGEs shape bacterial genome plasticity and facilitate the acquisition of traits such as antimicrobial resistance and metabolic capabilities(11, 12). Their distribution across bacterial genomes can therefore provide insights into patterns of gene exchange and ecological adaptation.

Bacterial defence systems are central to these interactions because they restrict the entry and persistence of foreign genetic material. CRISPR–Cas and restriction–modification systems recognise and target foreign nucleic acids, while abortive infection and signalling-based systems limit phage propagation through other mechanisms(13–15). Defence genes frequently occur together in genomic regions known as defence islands, and their repertoires vary across bacterial lineages and environments. In response, phages and other MGEs have evolved counter-defence mechanisms, including anti-CRISPR proteins, that overcome host protection. This reciprocal adaptation shapes both susceptibility to infection and opportunities for gene acquisition, making defence and counter-defence architectures potentially informative ecological signatures(16–18).

An emerging approach is therefore to evaluate microbial authenticity through evolutionary coherence: agreement among signatures of HGT, MGE activity and phage–host co-evolution(12, 19). Genuine human-or cancer-associated bacteria may exhibit evidence of ecological embedding through participation in MGE-mediated gene exchange and detectable interactions with bacteriophages. We hypothesised that coordinated patterns across these features could help distinguish human-associated bacteria from probable contaminants. Examining interconnected gene exchange networks, defence system architectures and phage–host linkages together could thus provide a biologically informed strategy for distinguishing signal from noise in cancer microbiome datasets. Their value lies in providing convergent evidence alongside conventional contamination assessments.

Understanding these relationships may also inform future approaches to microbiome modulation(20). The host specificity of bacteriophages offers opportunities to target selected bacteria, but infection outcomes depend partly on the balance between bacterial defence and phage counter-defence. Characterising these mechanisms could help identify barriers to phage targeting and guide experimental investigation of bacteria with potential clinical relevance(21, 22). In cancer-associated microbiomes, such applications first require a clearer understanding of the origin and biological significance of the microorganisms detected.

Here, we investigated whether bacteria–phage co-evolutionary signatures and MGE profiles provide complementary evidence for distinguishing human-associated bacteria from probable contaminants in cancer metagenomes. We analysed 514 pre-assembled metagenome-assembled genomes (MAGs) from the metastatic cancer dataset derived from the Hartwig Medical Foundation cohort, (2), together with a complementary collection of 785 high-quality MAGs derived from colorectal cancer faecal metagenomes (1). These collections represent distinct sampling contexts: metastatic tumour biopsies and colorectal cancer faecal samples. We integrate MGE profiling, phage–host predictions, defence system architectures and phage lifecycle characterisation to examine complementary aspects of bacterial ecology and evolution. By assessing these genomic signals together, we aimed to evaluate their potential to strengthen the interpretation of microbial signals in cancer metagenomes and identify features relevant to subsequent experimental investigation.

## 6. Methods

### Genomic dataset used in this study

To investigate ecological coherence within cancer-associated microbiomes, we analysed two publicly available metagenomic datasets. The primary dataset comprised 514 high- and medium-quality metagenome-assembled genomes (MAGs) recovered from metastatic tumour samples in the pan-cancer cohort of Battaglia et al. (2024), which integrates metagenomic, genomic, transcriptomic, and clinical data from 4,160 metastatic tumour biopsies. The MAGs were obtained from the study’s public Zenodo repository and represent tumour-associated microbial communities across multiple cancer types and anatomical sites(2). To provide an independent comparison from a well-characterised cancer-associated microbiome, we additionally analysed 60 colorectal cancer stool metagenomes from the German cohort (1), distributed across two European Nucleotide Archive (ENA) accessions (PRJEB27928 and PRJEB6070). Their data were released in two batches: 38 patients first published by Zeller et al. (2014; ENA PRJEB6070) (23) and 22 additional patients sequenced for Wirbel et al. (2019; ENA PRJEB27928) (1). Raw paired-end reads were downloaded from ENA, subjected to human read depletion and quality filtering, and subsequently assembled and binned, yielding 785 high-quality MAGs (completeness ≥90%, contamination ≤5%) from 55 samples containing at least one qualifying bin (Supplementary Methods). The German sub-cohort was selected instead of the full 768-sample meta-analysis to minimise technical heterogeneity, as samples were processed using consistent laboratory and sequencing protocols. The inclusion of both tumour-derived and stool-derived MAG collections enabled cross-context evaluation of microbial signatures, supporting discrimination of genuine human-associated taxa from potential environmental or methodological contaminants.

### Taxonomic Classification

Identification of MGEs and phage-host interactions was conducted using a comprehensive bioinformatic workflow applied to all MAGs (Figure 1). MAGs were screened for contaminant DNA and sample cross-contamination using two converging lines of evidence, namely genome-resolved taxonomy and predicted phage-host ecology. Taxonomic classification was carried out using GTDB-Tk (v2.3.0, database R214) (24), which served as the primary contaminant trigger. A MAG was considered a candidate contaminant when GTDB-tk placed it outside any phylum or genus with an established human association (Table S3), with the iPHoP (v1.4.2) ecological profile used as corroborating evidence (25). Each flagged MAG then received a final classification from a literature review of its genus or lineage (Table 1), which took precedence over the iPHoP profile where the two disagreed.

**Figure 1:**
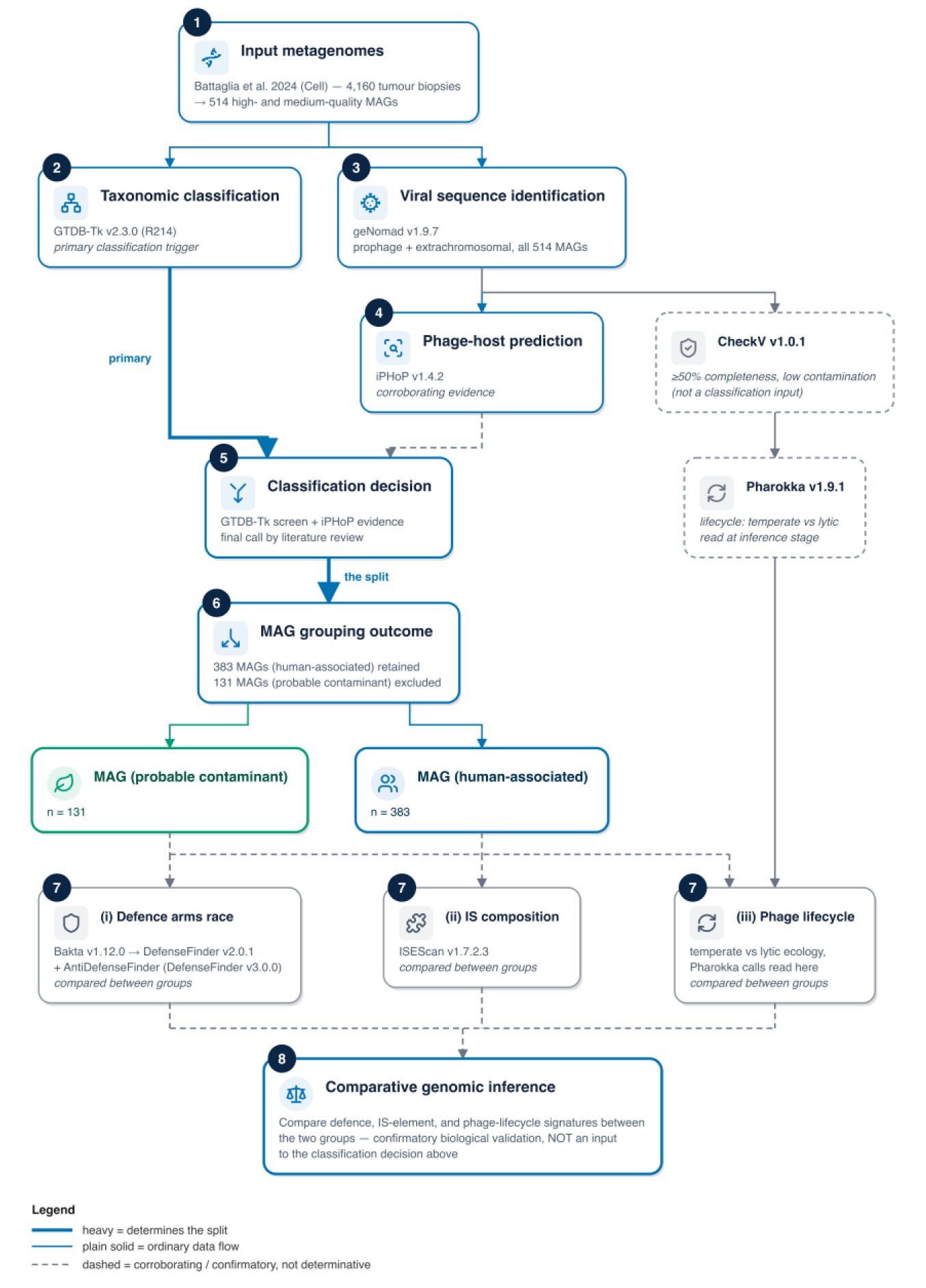
Study workflow: from input metagenomes to comparative genomic inference. **(1)** 514 high- and medium-quality metagenome-assembled genomes (MAGs) previously assembled and binned by Battaglia et al (2024) (2) was obtained from the study’s public Zenodo deposit. **(2-4)** Each MAG was taxonomically classified (GTDB-Tk v2.3.0, r214) and screened for viral sequence content (geNomad v1.9.7), whose output in turn fed phage-host prediction (iPHoP v1.4.2). **(5)** GTDB-Tk is the primary classification trigger (heavy edge, flagging any MAG placed outside a phylum or genus with established human association). iPHoP is corroborating evidence (dashed edge, flagging a MAG when ≥75% of its host predictions were environmental and 0% targeted a human-associated host). Flagged MAGs then receive a final call from a literature review of their genus or lineage, which takes precedence over iPHoP where the two disagree. **(6)** This produces the one determinative split of the whole workflow (heavy edge, labelled “the split”): 383 MAGs (human-associated) retained, 131 MAGs (probable contaminant) excluded. **(7)** The two resulting groups were then checked against three independent co-evolutionary signals, confirmatory biological validation applied after the 383/131 split rather than a classification criterion. The three signals are badged together as step 7, since they run in parallel rather than in sequence. (i) Defence systems arms race (Bakta v1.12.0 → DefenseFinder v2.0.1 + AntiDefenseFinder) and (ii) Insertion Sequence (IS) family composition (ISEScan v1.7.2.3) were both computed on the two classified groups and compared between them. (iii) Phage lifecycle ecology (temperate vs lytic) is analysed from the Pharokka (v1.9.1) call generated earlier in the workflow alongside CheckV filtered viral sequences. **(8)** All three signals converge on the same comparative genomic inference, confirmatory validation that the two groups are ecologically coherent. A similar analysis was also carried out for an independent cohort of 785 high-quality MAGs assembled from 55 colorectal cancer stool metagenomes (1), which is not shown in the figure. All 785 Wirbel MAGs were retained as human-associated at the classification step (Table 2) and are included as a comparison cohort in the subsequent figures.

**Table 1.** Genera classified as probable environmental contaminants at the GTDB-Tk screening stage.

| Genus | MAGs (n) | Order | Isolation source | Key references |
|---|---|---|---|---|
| *Ralstonia* spp. (incl. *R. pickettii*, n=43)<sup>‡</sup> | 72 | Burkholderiales | Water systems, soil, DNA extraction kits | (26-29) |
| *Bradyrhizobium* spp.<sup>§</sup> | 29 | Rhizobiales | Legume root nodules, agricultural soil | (30-32) |
| *Pelomonas* spp.<sup>§</sup> | 13 | Burkholderiales | Freshwater, aquatic habitat | (33, 34) |
| *Novosphingobium* spp.<sup>§</sup> | 5 | Sphingomonadales | Mangrove sediment, landfill leachate, pharmaceutical water | (35-37) |
| *Sphingomonas* spp.<sup>§</sup> | 4 | Sphingomonadales | Freshwater lakes, rice rhizosphere, soil biofilms | (38, 39) |
| *Pseudomonas_E* (*P. yamanorum*)<sup>†</sup> | 2 | Pseudomonadales | Subantarctic soil | (40, 41) |
| *Curvibacter* spp.<sup>§</sup> | 2 | Burkholderiales | River water, cyanobacterial bloom freshwater, polluted soil | (42, 43) |
| *Stagnimonas sp.* <sup>††</sup> | 1 | Nevskiales | Freshwater mesocosm | (44) |
| *Brachymonas* sp.<sup>§,\*\*</sup> | 1 | Burkholderiales | Petroleum-refinery wastewater, chironomid egg mass, soybean-processing waste sludge | (45-47) |
| Unclassified (no GTDB genus)<sup>¶</sup> | 2 | Caulobacterales | DNA extraction kits and laboratory reagents | (26) |

† Species confirmed by GTDB-Tk v2.3.0 (R214) assignment (exact match). Cited reference is for the same species.
‡ 43 MAGs assigned to *R. pickettii\_B* (GTDB subclade of *R. pickettii*). 29 assigned to *Ralstonia* sp001078575 (unnamed).
§ GTDB-Tk assigned these MAGs to unnamed species-level clades. Cited references support genus-level ecological classification only, not a confirmed species match.
¶ GTDB-Tk assigned no genus to these MAGs. Evidence is at order level: the cited reference lists three Caulobacterales genera among contaminants detected in sequenced negative controls, not the taxon itself.
\*\* GTDB's *Brachymonas* (45) is a later homonym of *Brachymonas* Grassé 1952. A replacement name, *Allobrachymonas*, has been proposed but is not yet reflected in the GTDB release used here.
†† GTDB-Tk assigned this MAG to genus only, with no species level match. The cited reference describes the genus.

**Table 2.**
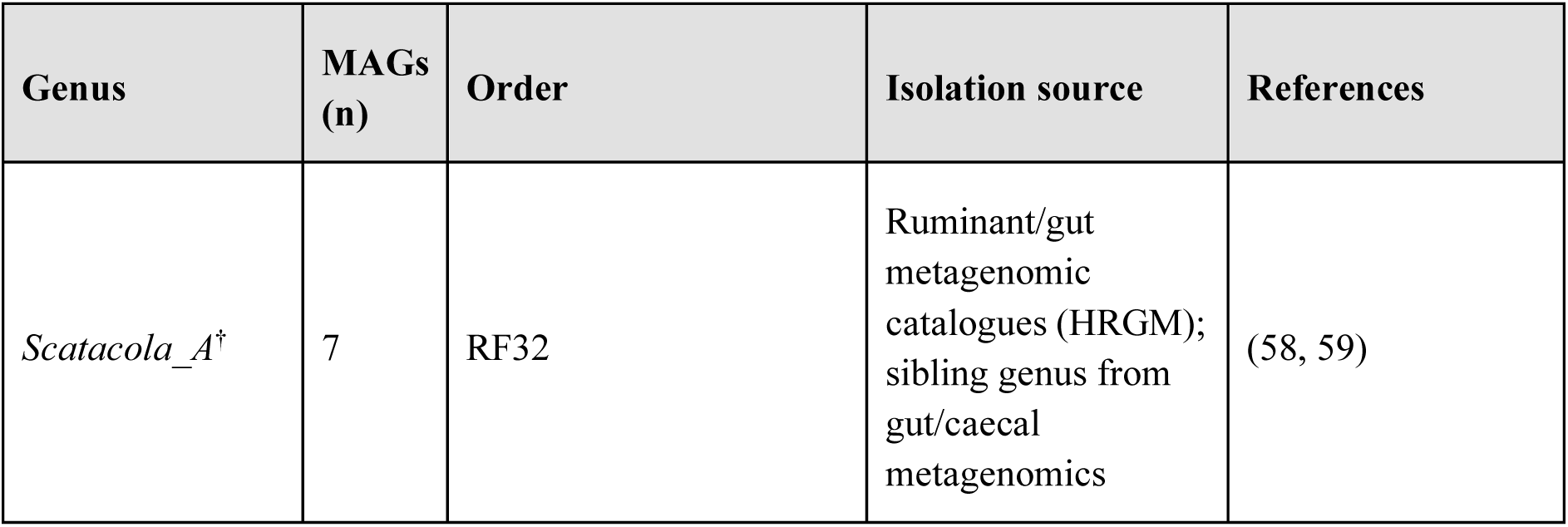
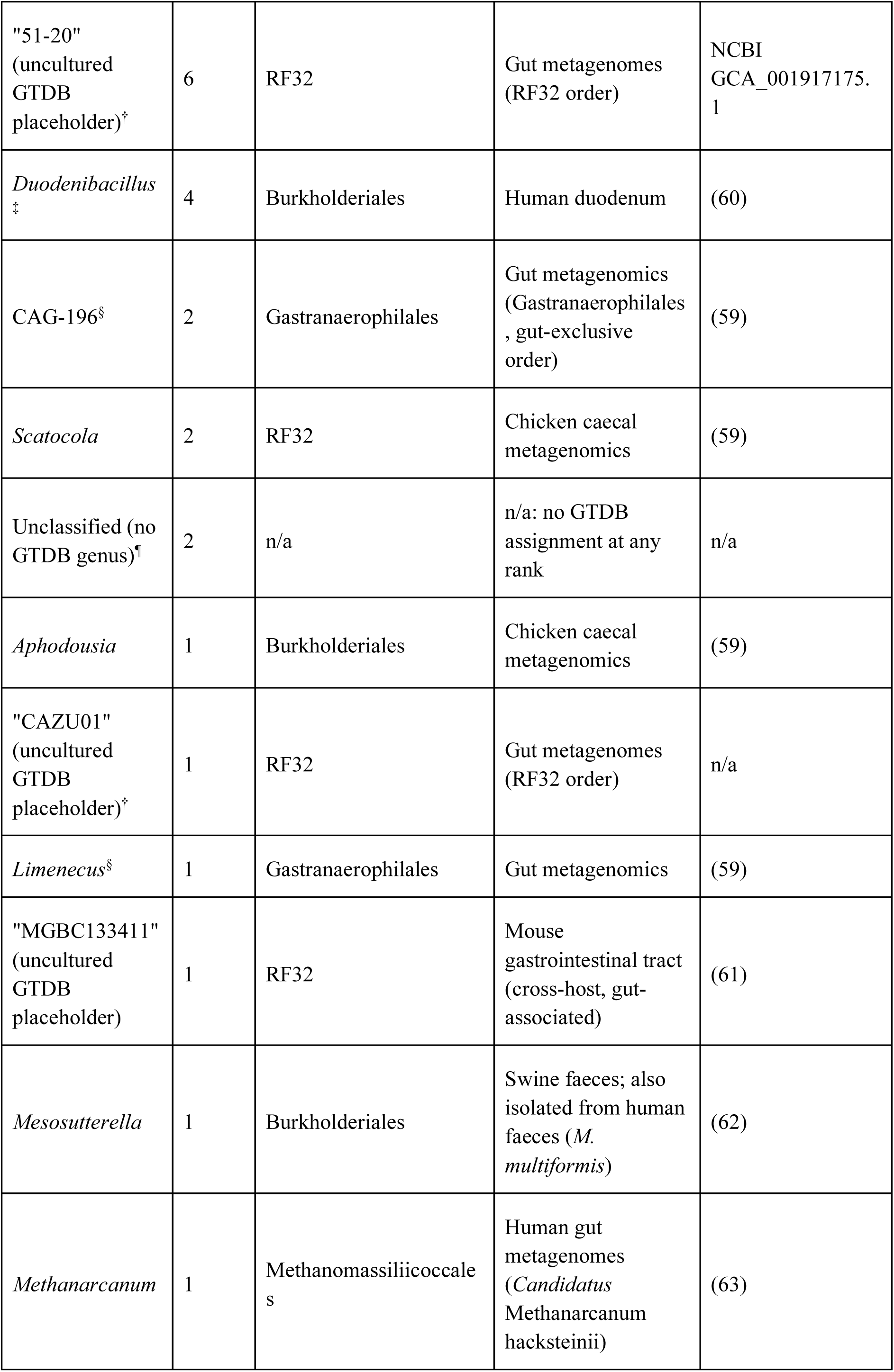
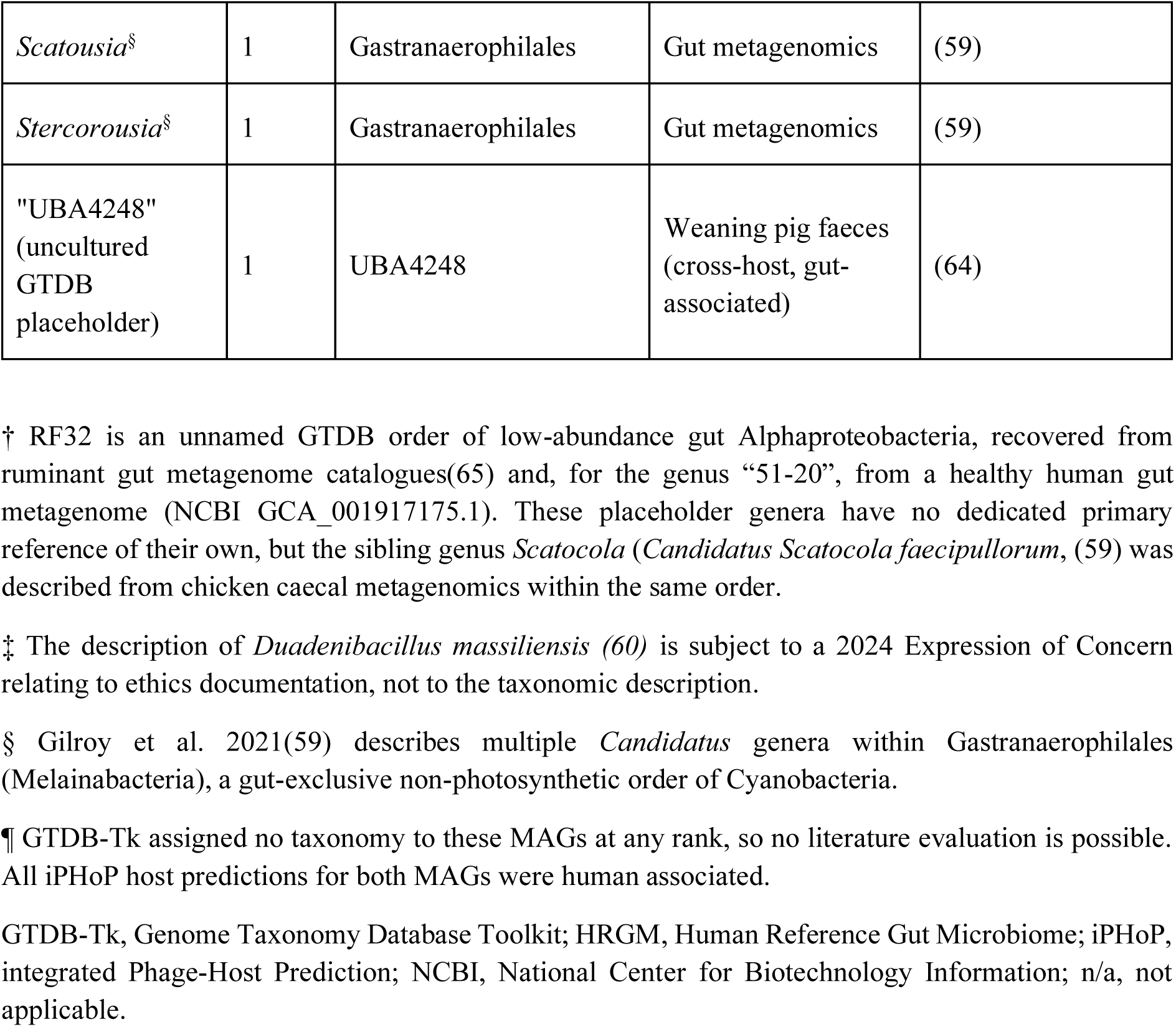
Genera of the 32 (1) metagenome-assembled genomes (MAGs) from the Wirbel colorectal cancer (CRC) cohort flagged by GTDB-Tk for unfamiliar or provisional genus level taxonomy. Thirty were confirmed as genuine gut commensals by literature review. The remaining two had no GTDB-Tk assignment at any rank and were retained because all their iPHOP host predictions were human-associated.

Viral sequences, including integrated prophages and extrachromosomal phage elements, were identified using geNomad (v1.9.7) (48), which integrates gene content, viral hallmark proteins and sequence-based classification. Plasmid sequences were identified in the same geNomad run. Across the 514 Hartwig MAGs, geNomad identified 2,902 viral sequences distributed among 431 MAGs. These sequences were retained without filtering by CheckV-estimated quality (v1.0.1) (49) to maximise sensitivity, including the recovery of fragmented viral sequences, and were processed through two complementary analytical pathways. For phage–host prediction, all geNomad-derived viral sequences were analysed using iPHoP (v1.4.2) (25). Predicted host lineages were reported using GTDB taxonomy at genus level. Hosts were assigned to human-associated or environmental categories using literature-curated phylum and genus lists (Table S3), with genus-level assignments capturing human-associated Proteobacteria such as *Neisseria*, *Haemophilus* and *Escherichia*. Host predictions were aggregated for each MAG to calculate the proportions assigned to each ecological category. A MAG received a contaminant flag when at least 75% of its predictions were assigned to environmental hosts and none were assigned to human-associated hosts. This study-specific rule was chosen to require a strong predominance of environmental affiliations without conflicting evidence of human association. It was distinct from iPHoP’s confidence threshold for individual host predictions (score ≥90), associated with an estimated genus-level false discovery rate below 10% in the published benchmarks(25). All flagged MAGs had exclusively environmental host predictions, indicating that their flags were not driven by values close to the 75% threshold. Ecological flags provided corroborating evidence, while final classification was determined by literature assessment of the assigned bacterial genus or lineage.

### Contaminant identification by using co-evolutionary signal integration

To robustly distinguish ecologically coherent human-associated microbes from potential contaminants, each MAG was evaluated against independent co-evolutionary signals: (i) phage–defence dynamics, (ii) insertion sequence (IS) element family composition, and (iii) phage lifecycle ecology.

Gene-level annotation was performed using Bakta (v1.12) (50, 51), which predicts open reading frames (ORFs), assigns functional annotations, and generates comprehensive GFF3 and protein FASTA (FAA) outputs for each MAG. The resulting FAA files were used as input for all subsequent defence-related tools. Defence and anti-defence systems were identified using DefenseFinder (v2.0.1). To characterise the phage-derived counter-defence layer, AntiDefenseFinder was used to detect phage-encoded proteins that antagonise bacterial immunity, thereby capturing an additional dimension of the bacteria–phage arms race (15, 52). Mobile genetic element features were characterised by using ISEScan (v1.7.2.3)(53).

### Analysis of phage life cycle

Viral sequences identified by geNomad were quality assessed using CheckV (v1.0.1) (49) and sequences meeting defined thresholds (≥50% completeness and low contamination) were retained for functional annotation using Pharokka (v1.9.1) enabling characterisation of phage lifecycle features, including integrase and excisionase genes indicative of temperate phages, as well as structural and lysis-associated proteins (54).

### Reference genome validation

To assess whether the defence and IS patterns observed in MAGs reflected lineage-associated features, MAGs were compared with complete reference genomes of the same genus. For the five most abundant genera in each of the Hartwig human-associated, Hartwig probable-contaminant and Wirbel MAG sets, all RefSeq assemblies at the Complete Genome level were downloaded from NCBI (July 2026; 3248 genomes). Each reference genome was classified as human or non-human in source from its BioSample host and isolation source fields, supplemented by a keyword search of BioSample and BioProject descriptions. Genomes of unresolved source were excluded. Reference genomes were annotated with Bakta, and defence systems and IS elements were identified in chromosomal replicons using DefenseFInder and ISEScan, as for the MAGs. Only genera with at least 10 genomes were analysed.

### Statistical analysis

All analyses were performed in R (v4.5.3) using the tidyverse (v2.0), with figures produced using ggplot2 (v4.0.2) and patchwork (v1.3.2), and statistical tests from the base stats package. Defence and anti-defence subtypes and IS families were scored as present and absent in each MAG, and prevalence was calculated as the percentage of MAGs in a group carrying at least one instance. Differences in prevalence between Hartwig human-associated MAGs and each comparison group (Hartwig probable contaminant, Wirbel MAGs, human and non-human reference genomes) were tested using two-sided Fisher’s exact tests. P values were corrected for multiple testing using the Benjamini-Hochberg (BH) method across all subtypes or families detected in at least one of the two groups compared, and adjusted P < 0.05 was considered significant. Per-MAG genome size, GC content, IS element, provirus and plasmid counts were compared between groups using two-sided Mann-Whitney U tests, and the proportion of temperate proviral contigs using Fisher’s exact tests, with BH correction across the three pairwise group comparisons in each analysis. The proportion of MAGs carrying at least one phage predicted to infect a human-associated host was compared between human-associated MAGs and probable contaminants using Fisher’s exact test. For reference genome validation, defence system and IS-element counts were normalised per Mbp of genome (chromosome only for reference genomes). MAGs were then compared with reference genomes of the same genus using Mann-Whitney U tests, BH-corrected across genera within each comparison. The association between genome size and IS element count was assessed using Spearman’s rank correlation.

## 7. Results

### Integrated taxonomic and habitat-based screening distinguishes human-associated MAGs from probable contaminants in cancer metagenomes

Cancer metagenomes typically contain low microbial biomass and are particularly susceptible to contamination from DNA extraction reagents and environmental sources (26, 37, 55). To address this, we screened 514 metagenome-assembled genomes (MAGs) using an integrated approach combining genome-resolved taxonomy, phage–host ecology, and literature-based habitat validation (Figure 1). Throughout this study, we refer to MAGs supported as members of the human microbiota as *Hartwig cancer-metagenome human-associated MAGs*, whereas MAGs with evidence of environmental or reagent origin are described as *Hartwig cancer-metagenome probable contaminant MAGs*. Concordance between MAG taxonomy and the predicted host affiliation of associated viral sequences was considered supportive of ecological coherence, whereas substantial discordance could indicate contamination, mis-binning, incidental co-assembly, or uncertainty in phage–host prediction.

GTDB-Tk identified 131 MAGs with no established affiliation with the human microbiota. Of these, 116 were independently supported by phage–host profiles dominated by environmentally associated taxa, whereas 14 yielded no iPHoP host predictions and therefore provided no ecological evidence contradicting their taxonomic classification (Table S1). One *Brachymonas* MAG contained a viral sequence for which iPHoP predicted *Neisseria*, a human-associated genus, as the host. This assignment was supported by only one prediction method and had low average amino-acid identity (AAI; 41.1%) and was therefore considered insufficient to override the taxonomic and habitat evidence. Such an isolated assignment is also consistent with the host-rounding behaviour reported for iPHoP, whereby poorly represented phages may be assigned to better-characterised related taxa(25). Literature-based habitat assessment instead supported an environmental origin for *Brachymonas*, as formally described species have been isolated from petroleum refinery wastewater plant, activated sludge, and chironomid egg masses rather than established human-associated environments(46, 56, 57). This MAG was therefore classified within the Hartwig cancer-metagenome probable contaminant group.

Overall, 131 of the 514 Hartwig MAGs (25.5%) were classified as probable contaminants. This group was strongly dominated by *Ralstonia* (n = 72; 55%) and *Bradyrhizobium* (n = 29; 22%), followed by *Pelomonas* (n = 13; 10%), *Novosphingobium* (n = 5; 4%), and *Sphingomonas* (n = 4; 3%). Less frequent genera shown in Figure 2a included *Curvibacter* and *Pseudomonas_E* (n = 2 each), and *Brachymonas, Stagnimonas* (n = 1 each), and two further MAGs with no formal genus assignment (order Caulobacterales, n = 2) (Table 1). Several of these genera are associated primarily with water, soil, plants, or laboratory environments, consistent with their classification as probable environmental or reagent-derived contaminants. Exclusion of these 131 MAGs resulted in a curated set of 383 Hartwig human-associated MAGs. Notably, no genera were shared between the human-associated and probable contaminant groups, supporting a clear taxonomic separation between the two classifications. The most abundant genera in this group were *Prevotella* (n = 37; 9.7%), *Fusobacterium* (n = 24; 6.3%), *Bacteroides* (n = 18; 4.7%), *Streptococcus* (n = 17; 4.4%), and *Leptotrichia* (tied with *Bulleidia at* n = 9; 2.3%) (Figure 2a).

**Figure 2:**
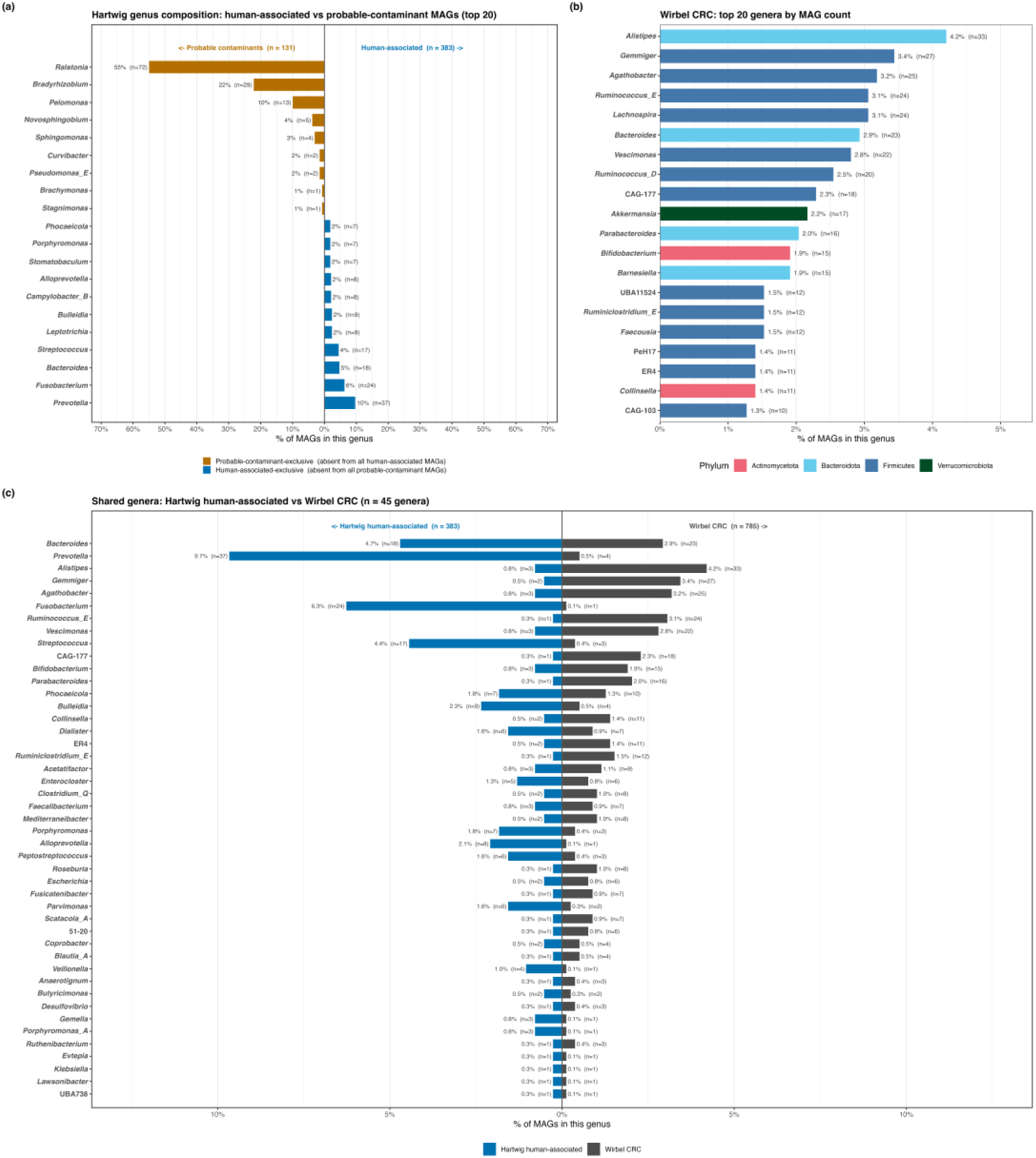
Genus composition of the Hartwig human-associated MAG set, the probable-contaminant set, and the Wirbel CRC cohort. **(a)** Genus composition of the 383 human-associated metagenome-assembled genomes (MAGs) versus the 131 probable-contaminant MAGs. Bars show the percentage of MAGs in each group assigned to a given genus, extending left for probable-contaminant MAGs (orange) and right for human-associated MAGs (blue). The panel shows the top 20 genera: every genus found among the probable-contaminant MAGs, plus the largest human-associated genera by MAG count. Of 134 genera present across the full 514-MAG set, none are shared between the human-associated and probable-contaminant groups. Two probable-contaminant MAGs lack a genus level assignment and are not plotted. **(b)** Genus composition of the Wirbel colorectal cancer (CRC) cohort (n = 785 MAGs), top 20 of 174 distinct genera by MAG count, coloured by phylum. **(c)** The 45 genera shared between the Hartwig human-associated set (125 genera) and the Wirbel CRC cohort (174 genera), all shown. Mirrored bars show the percentage of MAGs in each dataset assigned to that genus (blue = Hartwig human-associated, n = 383; dark grey = Wirbel CRC, n = 785), sorted by combined MAG count across both datasets.

To evaluate the performance of the same ecological screening approaches in an independent cancer dataset, we analysed 785 MAGs reconstructed from the colorectal cancer cohort of (1), hereafter referred to as *Wirbel 2019 CRC human-associated MAGs*. The Wirbel cohort showed a taxonomically broader gut-associated composition, with *Alistipes* (n = 33; 4.2%), *Gemmiger* (n = 27; 3.4%), *Agathobacter* (n = 25; 3.2%), *Ruminococcus_E* and *Lachnospira* (n = 24 each; 3.1%) (Figure 2b). Hartwig MAGs shared 45 genera with the independent Wirbel CRC cohort (Figure 2c). GTDB-Tk initially identified 32 MAGs with unfamiliar or provisional genus-level assignments. However, literature-based curation showed that 30 of the 32 belonged to recognised or emerging human-associated lineages rather than established environmental taxa (Table 2), while the remaining 2 lacked GTDB-Tk classification at any rank, so they could not be assessed against the literature, but their phage-host predictions pointed exclusively to human-associated hosts, and they were retained. No MAGs therefore met our criteria for probable environmental contamination in the Wirbel cohort (Table S2). These findings suggest that the combined screening method can distinguish poorly characterised human-associated bacteria from likely environmental or reagent-derived MAGs while reducing the risk of excluding genuine human-associated taxa.

### Conserved defence and anti-defence repertoires are consistent with ecological separation of human-associated MAGs from probable contaminants

To determine whether defence and anti-defence repertoires encode ecological signals indicative of host association, we compared Hartwig human-associated MAGs, Hartwig probable-contaminant MAGs, and the independent colorectal cancer (CRC) MAG collection from (1). Because interactions between bacteria, bacteriophages, and other MGEs drive reciprocal evolutionary adaptation, microbial genomes accumulate defence systems while MGEs evolve counter-defence mechanisms to overcome these barriers (18, 66, 67). The resulting defence architecture therefore represents a record of long-term ecological and evolutionary interactions. We tested whether human-associated MAGs exhibited defence and anti-defence signatures distinct from probable contaminants and whether these signatures were reproducible across an independent cancer microbiome dataset (Figure 3).

**Figure 3:**
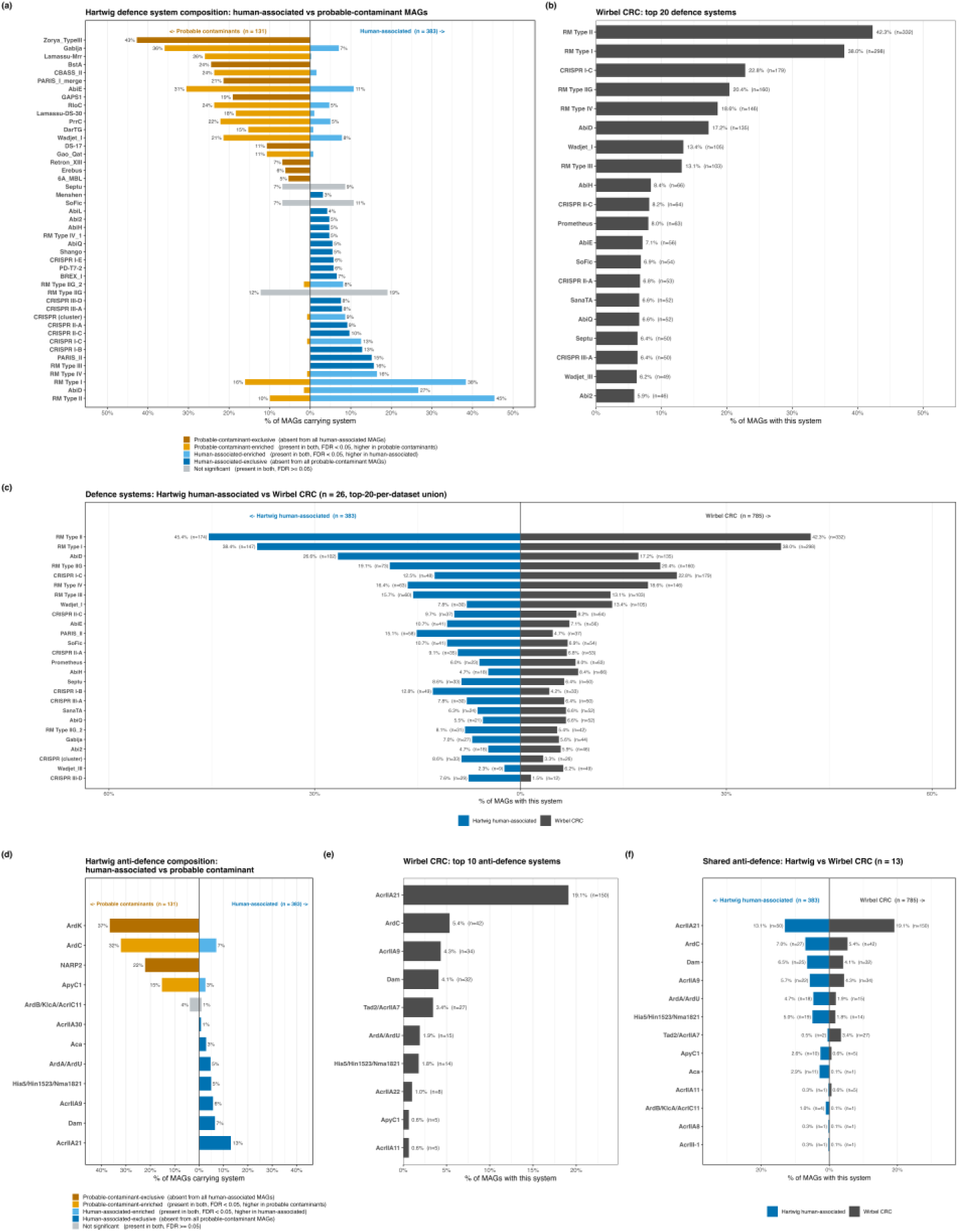
Defence and anti-defence system composition across all three datasets, and the shared core between Hartwig and Wirbel. **(a)** DefenseFinder defence subtype composition of the 383 human-associated metagenome-assembled genomes (MAGs) versus the 131 probable-contaminant MAGs. The panel shows the 20 most prevalent subtypes in each group, together with any subtype exclusive to one group and present at ≥3% prevalence (45 subtypes total). Bars show the percentage of MAGs in each group carrying a given subtype, extending left for probable-contaminant MAGs (orange) and right for human-associated MAGs (blue). Dark orange and dark blue indicate subtypes exclusive to one group (0% in the other). Orange and sky blue indicate significant enrichment in probable-contaminant and human-associated MAGs respectively, among subtypes present in both groups (Fisher’s exact test, Benjamini-Hochberg correction, false discovery rate (FDR) < 0.05). Subtypes present in both groups without a significant difference are shown in grey. **(b)** Defence composition of the Wirbel colorectal cancer (CRC) cohort (n = 785 MAGs), top 20 of 195 distinct DefenseFinder subtypes by prevalence. **(c)** Defence subtypes shared between the Hartwig human-associated MAG set and Wirbel CRC (n = 26 subtypes: the union of each dataset’s own top 20 by prevalence, 14 shared / 6 Hartwig-only / 6 Wirbel-only). Mirrored bars show the percentage of MAGs in each dataset (blue = Hartwig human-associated, n = 383; grey = Wirbel CRC, n = 785). **(d)** Anti-defence (ADF) system composition of human-associated versus probable-contaminant MAGs (12 subtypes: the union of the top 10 subtypes by prevalence in each group). Same colour and layout convention as panel (a). **(e)** ADF composition of the Wirbel CRC cohort (n = 785 MAGs), top 10 of 23 distinct ADF subtypes by prevalence. **(f)** 13 ADF subtypes shared between the Hartwig human-associated MAG set and Wirbel CRC, out of 23 distinct subtypes in each. Same mirrored bar convention as panel (c).

Restriction–modification (RM) systems were among the most prevalent defence systems in both cohorts (Figure 3a-c). RM type II occurred in 45.4% of Hartwig MAGs and 42.3% of Wirbel MAGs, while RM type I occurred in 38.4% and 38.0%, respectively. Other RM systems were also represented at comparable frequencies, including RM type IIG (19.1% versus 20.4%), RM type IV (16.4% versus 18.6%), and RM type III (15.7% versus 13.1%) in Hartwig and Wirbel, respectively. Both cohorts additionally contained CRISPR–Cas, abortive-infection and other accessory defence systems, although their relative prevalence varied. AbiD (26.6% versus 17.2%), PARIS-II (15.1% versus 4.7%), CRISPR I-B (12.8% versus 4.2%), SoFic (10.7% versus 6.9%) and CRISPR III-D (7.6% versus 1.5%) were more frequent in Hartwig human-associated MAGs, whereas CRISPR I-C (12.5% versus 22.8%), Wadjet-I (7.8% versus 13.4%) and AbiH (4.7% versus 8.4%) were more prevalent in the Wirbel cohort. Across the 26-system union of the top 20 defence profiles from the two datasets, substantial overlap was observed (14 ranked in the top 20 of both), indicating conservation of major defence-system classes across independently reconstructed cancer-associated microbiomes despite differences in the prevalence of individual systems (Table S4).

In contrast, Hartwig probable-contaminant MAGs showed a markedly different defence composition (Figure 3a). Several systems were particularly prominent in this group, including Zorya type III (43%), Gabija (36%), AbiE (31%), Lamassu-Mrr (26%), BstA (24%), CBASS type II (24%), RloC (24%), PrrC (22%), PARIS-I_merge (21%), and Wadjet-I (21%) (all BH-adjusted P < 0.05). Several of these systems were absent (Zorya type III, BstA, PARIS-I_merge) or occurred only at low prevalence (Lamassu-Mrr, RloC, PrrC, CBASS type II) among the human-associated MAGs. Thus, defence-system composition provided an ecological feature differentiating the two Hartwig MAG groups.

We next examined anti-defence determinants as an additional indicator of historical interactions with phages and other MGEs. Consistent with the high prevalence of restriction-modification (RM) and CRISPR-Cas systems among human-associated MAGs, several anti-RM and anti-CRISPR determinants were detected (Figure 3d-f). This coexistence is evolutionarily plausible, as recurrent interactions between bacteria and invading MGEs can drive the parallel accumulation of defence systems and corresponding counter-defence mechanisms. Anti-CRISPR genes are frequently associated with other anti-defence functions, including anti-restriction proteins such as ArdA and KlcA, which can enhance MGE establishment and dissemination by antagonising bacterial RM systems(18). Thirteen anti-defence subtypes were shared between the Hartwig human-associated and Wirbel CRC datasets (Figure 3f). Anti-CRISPR A*crIIA21* was the most prevalent shared subtype in both cohorts, occurring in 13.1% (50/383) of Hartwig human-associated MAGs and 19.1% (150/785) of Wirbel MAGs. Other shared anti-defence determinants included ArdC (7.0% versus 5.4%), AcrIIA9 (5.7% versus 4.3%), Dam (6.5% versus 4.1%), Arda/ardu (4.7% versus 1.9%), and Hia5/hin1523/nma1821 (5.0% versus 1.8%). The probable-contaminant group again showed a different profile (Figure 3d; Table S5). ArdK (36.6%) and ArdC (32.1%) were the most prevalent anti-defence subtypes among probable contaminants, followed by NARP2 (22.1%) and ApyC1 (15.3%). ArdK and NARP2 were not detected in human-associated MAGs, and ArdC and ApyC1 were less frequent there (7.0% and 2.6%; all BH-adjusted P < 0.001). In contrast, several of the more prevalent Hartwig human-associated anti-defence subtypes, including AcrIIA21, Dam, AcrIIA9, Hia5, ArdA and Aca, were not detected in the probable-contaminant MAGs.

Together, the conservation of major defence and anti-defence features between the Hartwig human-associated and independent Wirbel CRC datasets, alongside their contrasting distribution in the probable-contaminant MAGs, provides additional ecological evidence supporting the distinction between human-associated microbial genomes and probable environmental or reagent-derived contaminants.

Several defence systems enriched in probable-contaminant MAGs, including Gabija and Zorya type III, lacked corresponding anti-defence annotations. Although phage-encoded inhibitors have been reported for Gabija and other recently characterised defence systems(68), their absence in the current dataset may reflect incomplete recovery of MGE sequences within MAGs, sequence divergence from currently characterised inhibitors, genuine absence of cognate counter-defence genes, or limitations of existing annotation databases. Because AntiDefenseFinder relies on reference models derived from experimentally characterised families(52), highly divergent or previously undescribed anti-defence proteins are unlikely to be detected.

Consequently, establishing specific bacteria-phage arms-race relationships will require analyses beyond cohort-level prevalence patterns. Future investigations should incorporate MAG-level co-occurrence analyses of anti-defence determinants within prophages, plasmids, and other MGEs, and functional validation of predicted defence-counter-defence pairs to determine whether particular anti-defence proteins are directly associated with their cognate host defence systems.

### Insertion sequence repertoires differentiate human-associated and probable contaminant MAGs

To determine whether mobile-element composition provided an additional genomic signal for distinguishing human-associated microbial genomes from probable contaminants, we compared insertion sequence (IS) family composition between Hartwig cancer-metagenome human-associated MAGs and Hartwig probable-contaminant MAGs using ISEScan (53). Probable-contaminant MAGs carried a significantly greater IS burden than human-associated MAGs (median 12 versus 6 IS elements per MAG; Mann-Whitney P = 2.9 × 10⁻¹⁷), indicating substantial differences in mobile-element content between the two groups.

IS-family composition further accentuated this distinction. Of the 25 IS families detected, 19 differed significantly in prevalence after multiple-testing correction (Fisher’s exact test, Benjamini-Hochberg FDR < 0.05; Figure 4a; Table S6). Eleven families were enriched in probable-contaminant MAGs, whereas eight were enriched in human-associated MAGs. Among the latter, IS200/IS605 showed the strongest enrichment, occurring in 36.6% (140/383) of human-associated MAGs compared with 1.5% (2/131) of probable-contaminant MAGs. IS982 (12.8%, 49/383), IS1634 (7.8%, 30/383), and IS607 (4.7%, 18/383) were detected exclusively in human-associated MAGs, while additional enrichment was observed for ISL3, IS1182, IS4, and ISNCY. In contrast, probable-contaminant MAGs exhibited marked enrichment of IS66 (76.3%, 100/131 versus 13.3%, 51/383), IS1595 (44.3%, 58/131 versus 7.3%, 28/383), and unclassified (“new”) IS elements (54.2%, 71/131 versus 1.0%, 4/383). Additional contaminant-enriched families included IS3, IS5, IS630, IS21, IS256, IS110, IS481, and IS6. Collectively, these findings demonstrate that the Hartwig human-associated and probable-contaminant MAG groups differ not only in overall IS burden but also in the structure of their insertion-sequence repertoires, supporting ecological differentiation between the two microbial genome collections.

**Figure 4:**
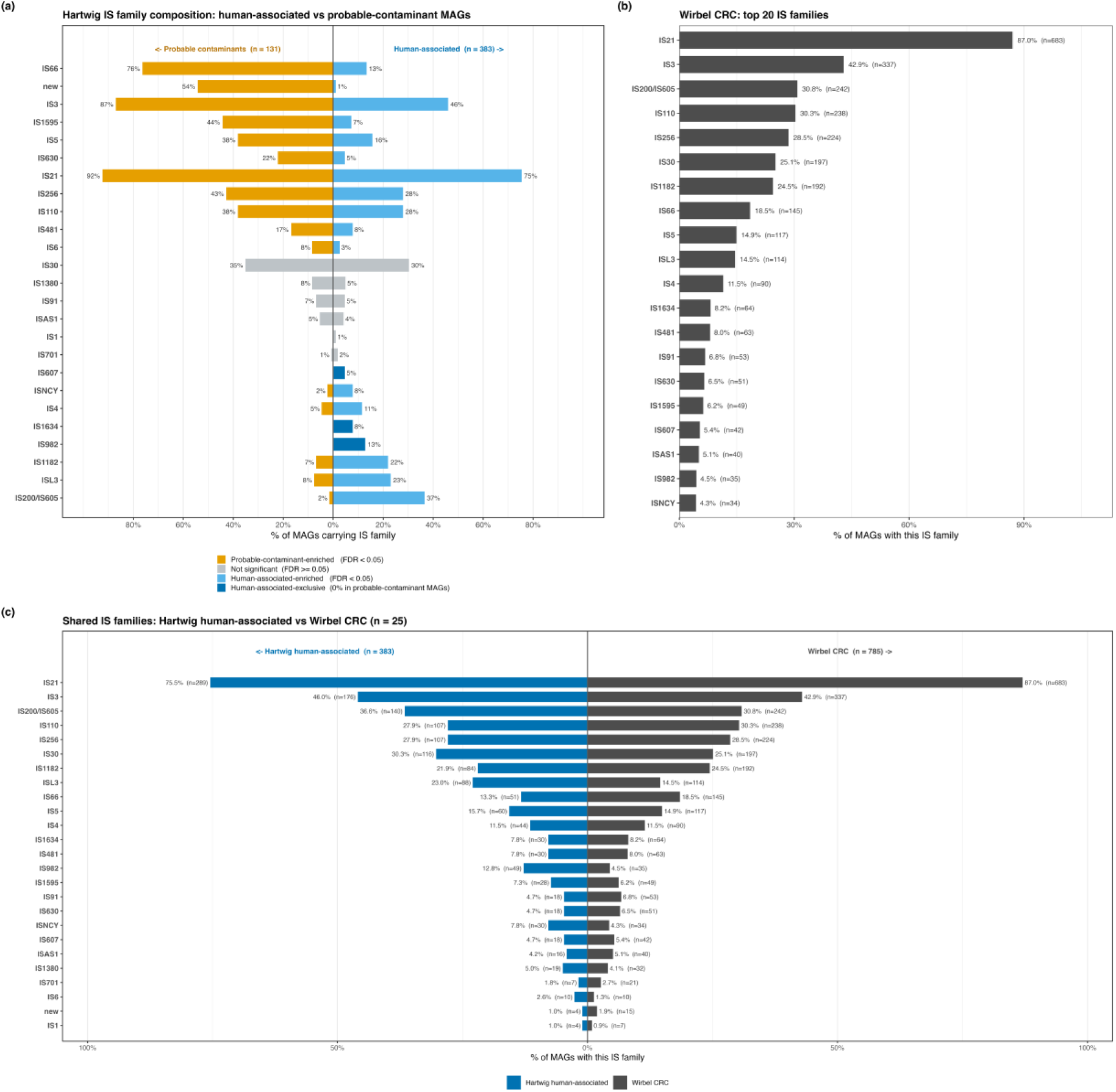
IS element family composition across all three datasets, and the shared core between Hartwig and Wirbel. **(a)** ISEScan IS family composition of the 383 human-associated metagenome-assembled genomes (MAGs) versus the 131 probable-contaminant MAGs. Bars show the percentage of MAGs in each group carrying a given IS family, extending left for probable-contaminant MAGs (orange) and right for human-associated MAGs (blue). All 25 IS families identified across the full 514-MAG set are shown. Human-associated MAGs carried all 25 families, whereas probable-contaminant MAGs carried 21. Each family was tested with Fisher’s exact test, and P values were corrected using the Benjamini-Hochberg method (false discovery rate, FDR). Dark blue indicates IS families exclusive to human-associated MAGs (0% in probable-contaminant MAGs; FDR < 0.05), orange indicates families enriched in probable-contaminant MAGs (FDR < 0.05), sky blue indicates families enriched in human-associated MAGs (FDR < 0.05), and non-significant results are shown in grey. **(b)** IS family composition of the Wirbel colorectal cancer (CRC) cohort (n = 785 MAGs), top 20 of 25 distinct ISEScan families by prevalence. **(c)** IS families shared between the Hartwig human-associated set and Wirbel CRC. All 25 families are present in both datasets, and all are shown. Mirrored bars show the percentage of MAGs in each dataset assigned to that family (blue = Hartwig human-associated, n = 383; grey = Wirbel CRC, n = 785), sorted by combined MAG count.

To assess whether the human associated IS signature was reproducible in the independent cancer microbiome dataset, we also analysed 785 MAGs reconstructed from the Wirbel CRC cohort. The overall IS burden closely resembled that observed in Hartwig human-associated MAGs (median 5 versus 6 IS elements per MAG) and remained substantially lower than that of Hartwig probable-contaminant MAGs. Notably, all 25 IS families identified in the Hartwig comparison were also detected in the Wirbel dataset (Figure 4c). Several predominant IS families exhibited comparable prevalence across cohorts, including IS21 (75.5% versus 87.0%), IS3 (46.0% versus 42.9%), IS200/IS605 (36.6% versus 30.8%), IS110 (27.9% versus 30.3%), and IS256 (27.9% versus 28.5%) (Figure 4b, c), indicating broad similarity in IS-family composition despite differences in sample origin and cohort structure.

Importantly, several IS families enriched in Hartwig human-associated MAGs were also recovered in the Wirbel cohort. IS982, IS1634, and IS607, which were absent from Hartwig probable-contaminant MAGs, were detected in 4.5% (35/785), 8.2% (64/785), and 5.4% (42/785) of Wirbel MAGs, respectively (Figure 4c). Additional human-associated-enriched families showed similar representation across cohorts, including IS1182 (21.9% versus 24.5%), IS4 (11.5% versus 11.5%), and ISNCY (7.8% versus 4.3%). Although ISL3 occurred at a lower frequency in Wirbel MAGs (14.5%) than in Hartwig human-associated MAGs (23.0%), it remained detectable in both datasets. These observations suggest that several IS-family patterns identified in the Hartwig human-associated MAGs are reproducible in an independent cancer-associated microbiome dataset.

In contrast, the contaminant associated IS signature identified in the Hartwig dataset was not observed to the same extent in the Wirbel cohort (Figure 4a, c). Unclassified (“new”) IS elements remained rare in Wirbel MAGs (1.9%, 15/785), resembling the Hartwig human-associated group (1.0%, 4/383) and contrasting with Hartwig probable contaminants (54.2%, 71/131). Similarly, IS66 occurred in 18.5% (145/785) of Wirbel MAGs and 13.3% (51/383) of Hartwig human-associated MAGs, compared with 76.3% (100/131) of probable-contaminant MAGs. IS1595 showed a comparable pattern, occurring in 6.2% (49/785) of Wirbel MAGs and 7.3% (28/383) of Hartwig human-associated MAGs, but in 44.3% (58/131) of probable contaminants.

Taken together, these results indicate that several IS-family features associated with the Hartwig human-associated MAGs are also present in an independent CRC-derived MAG collection, whereas the most prominent contaminant-associated signatures are not similarly represented.

### Divergent genome architecture characterises human-associated and probable-contaminant MAGs

Hartwig probable-contaminant MAGs were substantially larger than Hartwig human-associated MAGs, with median genome sizes of 5.0 and 2.0 Mbp, respectively (Mann-Whitney P = 7.2 × 10⁻^53^). Wirbel CRC MAGs exhibited an intermediate median genome size of 2.41 Mbp and therefore more closely resembled the Hartwig human-associated group (Tables S1 and S2). Larger bacterial genomes commonly encode broader metabolic, regulatory, and environmental-response capacities and can accommodate greater amounts of accessory DNA (Konstantinidis & Tiedje, 2004). Consistent with this relationship, IS burden correlated positively with genome size within both Hartwig human-associated MAGs (Spearman’s ρ = 0.434, P = 4.9 × 10⁻¹⁹) and probable-contaminant MAGs (ρ = 0.415, P = 8.4 × 10⁻⁷). When normalised to genome size, IS density was in fact lower in probable-contaminant MAGs than in human-associated MAGs (median 2.0 versus 2.8 IS elements per Mbp; Mann-Whitney P = 0.005), indicating that the greater IS burden of probable-contaminant MAGs reflects their larger genomes rather than a higher density of IS elements.

GC content provided an additional distinction between the two MAG groups. Hartwig human-associated MAGs had a median GC content of 43.0%, compared with 64.0% among probable-contaminant MAGs (Mann-Whitney P = 1.1 × 10⁻⁶¹). As GC content is strongly influenced by phylogeny and can covary with genome size and ecological strategy, these differences suggest contrasting taxonomic and evolutionary compositions between the two groups (69). Consistent with this interpretation, Wirbel CRC MAGs displayed an intermediate median GC content of 48.7%, despite genome sizes more closely resembling those of the Hartwig human-associated MAGs. Free-living environmental bacteria often possess larger, gene-rich genomes that facilitate metabolic versatility and adaptation to fluctuating environmental conditions (70), whereas adaptation to host-associated niches can relax selection on genes required for environmental survival and lead to genome reduction (71).

These analyses indicate that human-associated and probable-contaminant MAGs differ not only in mobile-element composition but also in broader aspects of genome architecture.

### Human-associated microbiomes exhibit temperate prophage enrichment and reduced plasmid burden

To further evaluate whether mobile genetic elements provided additional ecological features for distinguishing human-associated MAGs from probable contaminants, we compared phage-host associations, viral content, plasmid burden, and prophage characteristics across the dataset.

The iPHoP-based phage-host analysis showed strong concordance with MAG classification (Figure S1). Among the 383 Hartwig human-associated MAGs, 295 (77%) contained phages predicted to infect human-associated bacterial taxa, whereas 88 (23%) yielded no host prediction. In contrast, 116 of 131 probable-contaminant MAGs (88.5%) contained phages assigned to non-human-associated hosts, 14 (10.7%) yielded no prediction, and only a single MAG received a human-associated host assignment. The distribution of host predictions differed significantly between the two groups (Fisher’s exact OR = 430, P = 5.7 × 10⁻⁶¹), indicating strong agreement between bacterial taxonomy and phage-host ecological affiliation.

Overall viral burden showed a modest but statistically significant difference between groups. geNomad identified 2,017 viral sequences across Hartwig human-associated MAGs (5.3 per MAG; median 3) and 885 viral sequences across probable-contaminant MAGs (6.8 per MAG; median 5; Mann-Whitney P = 6.2 × 10⁻⁴). CheckV identified 139 proviral sequences within human-associated MAGs (0.36 per MAG) and 73 within probable-contaminant MAGs (0.56 per MAG) (Figure 5a). The Wirbel CRC MAGs contained 3,789 viral sequences in total (4.8 per MAG), a rate slightly below both Hartwig groups, and 446 CheckV-confirmed prophages (0.57 per MAG), closely matching the rate observed among Hartwig probable-contaminant MAGs.

**Figure 5:**
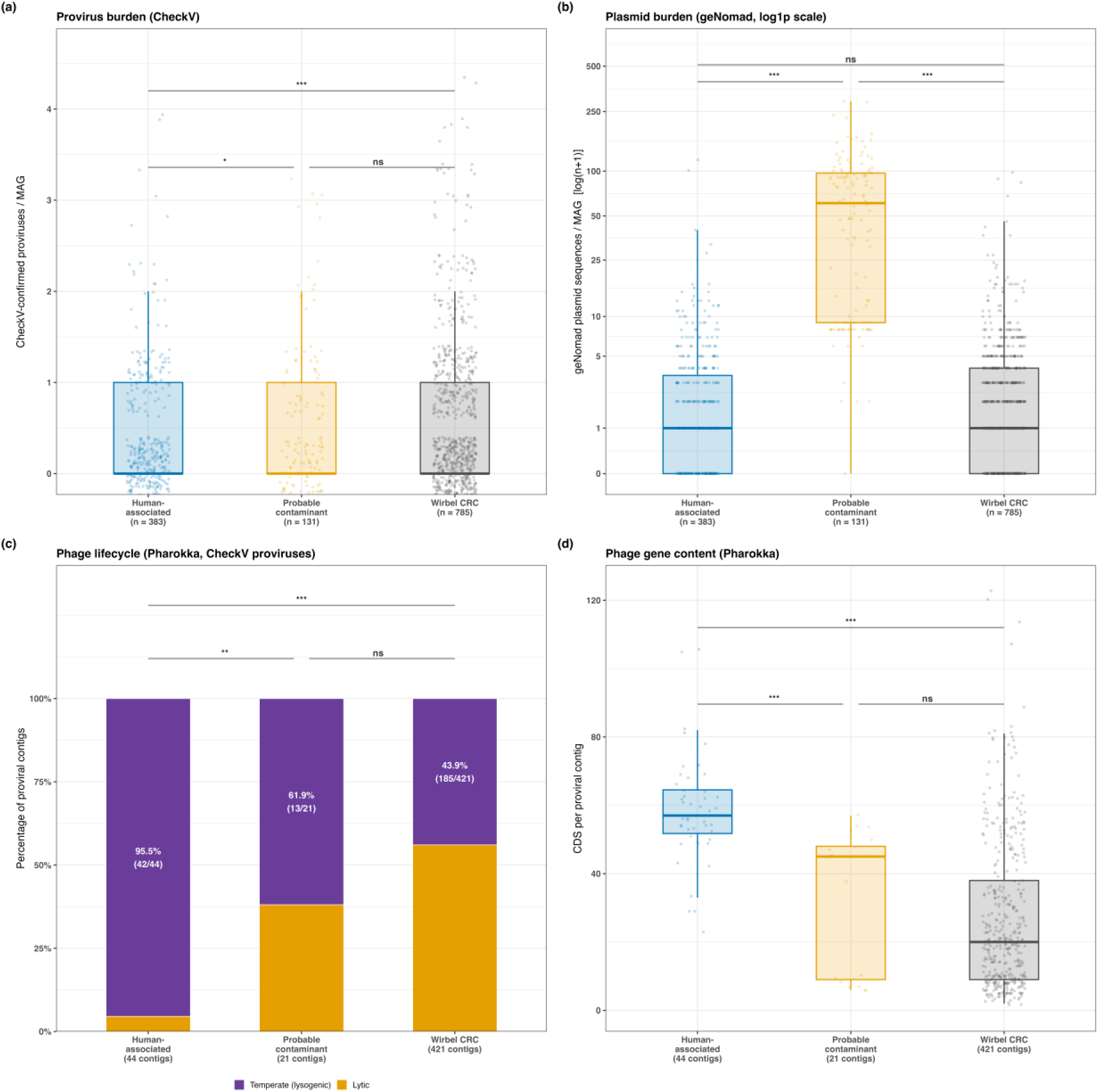
Mobile element landscape: proviruses, plasmids, and phage lifestyle across all three datasets. Panels (a) and (b) show human-associated (n = 383), probable-contaminant (n = 131) and Wirbel colorectal cancer (CRC, n = 785) metagenome-assembled genomes (MAGs). Panels (c) and (d) are restricted to CheckV-confirmed proviral contigs with a Pharokka annotation (44, 21 and 421 contigs for the same three groups). **(a)** CheckV-confirmed provirus burden per MAG (Mann-Whitney test). Human-associated MAGs had significantly lower provirus burden than probable-contaminant MAGs (p = 0.013), and Wirbel MAGs (p = 1.5 × 10^-4^). Probable-contaminant and Wirbel MAGs did not differ significantly. **(b)** geNomad plasmid burden per MAG, shown on a log (n + 1) scale (Mann-Whitney test). Probable-contaminant MAGs carried substantially more plasmid sequences than either other group (both p < 0.001). Human-associated and Wirbel MAGs did not differ significantly. **(c)** Phage lifecycle composition of proviral contigs classified from their Pharokka annotation as temperate (lysogenic) or lytic (Fisher’s exact test). Values in each bar give the percentage of temperate contigs, with the number of contigs over the total in brackets. Human-associated MAGs showed significantly higher odds of temperate proviruses than probable-contaminant MAGs (odds ratio (OR) = 12.3, *p* = 0.0017) and Wirbel MAGs (OR = 26.7, *p* = 1.2 × 10⁻^11^). Probable-contaminant and Wirbel MAGs did not differ significantly (OR = 2.1, *p* = 0.12). **(d)** Number of Pharokka-predicted coding sequences (CDSs) per proviral contig (Mann-Whitney test). Human-associated proviral contigs contained significantly more CDSs than those from probable-contaminant MAGs (*p* = 4.4 × 10⁻⁶) and Wirbel MAGs (*p* = 1.7 × 10⁻^15^). Probable-contaminant and Wirbel contigs did not differ significantly (*p* = 0.15). Boxplots in (a), (b) and (d) show the median and interquartile range (IQR), with 1.5 × IQR whiskers, and points show individual MAGs (a, b) or proviral contigs (d). P values are Benjamini-Hichberg corrected across the three pairwise comparisons within each panel.

In contrast, plasmid-associated sequence burden differed markedly between the two Hartwig groups (Figure 5b). geNomad detected 1,216 plasmid-associated sequences across Hartwig human-associated MAGs, corresponding to 3.2 sequences per MAG, compared with 8,677 sequences across probable-contaminant MAGs, corresponding to 66.2 sequences per MAG. This approximately 20-fold difference contrasted with the independent Wirbel cohort, in which 2,499 plasmid-associated sequences were detected across 785 MAGs (3.2 per MAG), closely matching the Hartwig human-associated average and remaining substantially lower than that observed among probable contaminants. The exceptionally larger plasmid burden among probable contaminants is consistent with the genomic characteristics of several dominant taxa in this group. Some *Ralstonia* and related environmental Proteobacteria possess multipartite genomes containing large accessory replicons, including chromids and megaplasmids (71, 72).

We next examined prophage-associated lifestyle features using Pharokka (Figure 5c). Integration- and excision-associated genes were detected in 95.5% (42/44) of Hartwig human-associated proviral contigs, compared with 61.9% (13/21) of probable-contaminant proviral contigs (Fisher’s exact OR = 12.32, P = 1.7× 10⁻^3^). These features were identified in 43.9% (185/421) of Wirbel CRC proviral contigs. Thus, proviruses recovered from Hartwig human-associated MAGs were more frequently associated with genetic features consistent with temperate lifestyles than those recovered from probable-contaminant MAGs.

Additional differences were observed in proviral gene content (Figure 5d). Pharokka-annotated proviral contigs from Hartwig human-associated MAGs encoded a median of approximately 57 CDS per contig, compared with 45 CDS among probable-contaminant proviruses and 20 CDS among Wirbel CRC proviruses. The difference between Hartwig human-associated and probable-contaminant proviruses was significant (Mann-Whitney P = 4.4 × 10⁻^6^), while the difference between Hartwig human-associated and Wirbel proviruses was more pronounced (P = 1.7× 10⁻^15^).

Taken together, these results indicate that mobile-element characteristics differ between Hartwig human-associated and probable-contaminant MAGs. While overall viral burden was broadly comparable across groups, human-associated MAGs were characterised by lower plasmid-associated sequence burden, stronger concordance with human-associated phage hosts, and a higher prevalence of prophage features associated with temperate lifestyles. Several of these characteristics, particularly the low plasmid burden, were also observed in the independent Wirbel CRC cohort, whereas the elevated plasmid-associated signal was identified in probable-contaminant MAGs.

### Reference-genome comparisons indicate predominantly lineage-associated defence and IS repertoires in human-associated MAGs

Several genera enriched among the Hartwig human-associated MAGs, including *Prevotella*, *Fusobacterium*, *Bacteroides*, and *Streptococcus*, are well-established members of the human microbiota(20, 73, 74). To assess whether the defence-system and IS patterns observed across the Hartwig and Wirbel cohorts primarily reflected lineage-associated genomic features, we compared representative MAGs with genus-matched reference genomes. The analysis included four dominant human-associated genera representing 96 of 383 Hartwig human-associated MAGs, three dominant probable-contaminant genera representing 114 of 131 probable-contaminant MAGs, and *Alistipes* from the independent Wirbel CRC cohort.

Across both human-associated and probable-contaminant lineages, MAGs generally exhibited lower defence-system and IS densities than their corresponding complete reference genomes. For example, *Prevotella* (2.75 versus 4.44 systems/Mbp) and *Bacteroides* (1.62 versus 2.18 systems/Mbp) MAGs contained fewer defence systems than matched human-source reference genomes (Figure S2a), while similar reductions were observed in environmental lineages such as *Ralstonia* (0.89 versus 1.83 systems/Mbp) and *Bradyrhizobium* (0.55 versus 1.30 systems/Mbp) (Figure S2c). A comparable trend was observed for IS density, with marked reductions relative to reference genomes in both human-associated genera, including *Prevotella* (2.69 versus 13.87 IS elements/Mbp) and *Fusobacterium* (3.95 versus 8.10 IS elements/Mbp) (Figure S2b), and environmental genera such as *Ralstonia* (1.97 versus 15.29 IS elements/Mbp) (Figure S2d). The same pattern was independently reproduced in Wirbel *Alistipes* MAGs (Figure S2e, f). Because these reductions occurred across diverse bacterial lineages and ecological groups, they are likely attributable to incomplete recovery of repetitive and accessory genomic regions during metagenomic assembly and binning.

We next examined whether the dominant defence systems and IS families identified in the human-associated MAGs were represented within lineage-matched reference genomes. The major defence signatures were largely conserved: all ten dominant defence systems identified in *Prevotella*, *Fusobacterium*, *Bacteroides*, and *Streptococcus* MAGs were also detected in corresponding reference genomes (Figure 6a-d), while nine of the ten dominant systems observed in Wirbel *Alistipes* MAGs were similarly recovered (Figure 6e). Representative lineage-associated signatures included PARIS-II and restriction-modification systems in *Prevotella* and *Bacteroides*; CRISPR I-B, BREX-I, AbiL, and PD-T7-2 in *Fusobacterium*; and multiple restriction-modification and CRISPR-Cas systems in *Streptococcus*. Similar patterns were observed for IS families. IS982, IS1634, and IS607, although absent from Hartwig probable-contaminant MAGs, were present in reference genomes, while the human-associated-enriched IS200/IS605 family was prominent among *Fusobacterium* references (Figure 6f-j). Together, these findings suggest that much of the defence-system and IS-family concordance observed between the Hartwig and Wirbel datasets reflects conserved lineage-associated genomic features.

**Figure 6:**
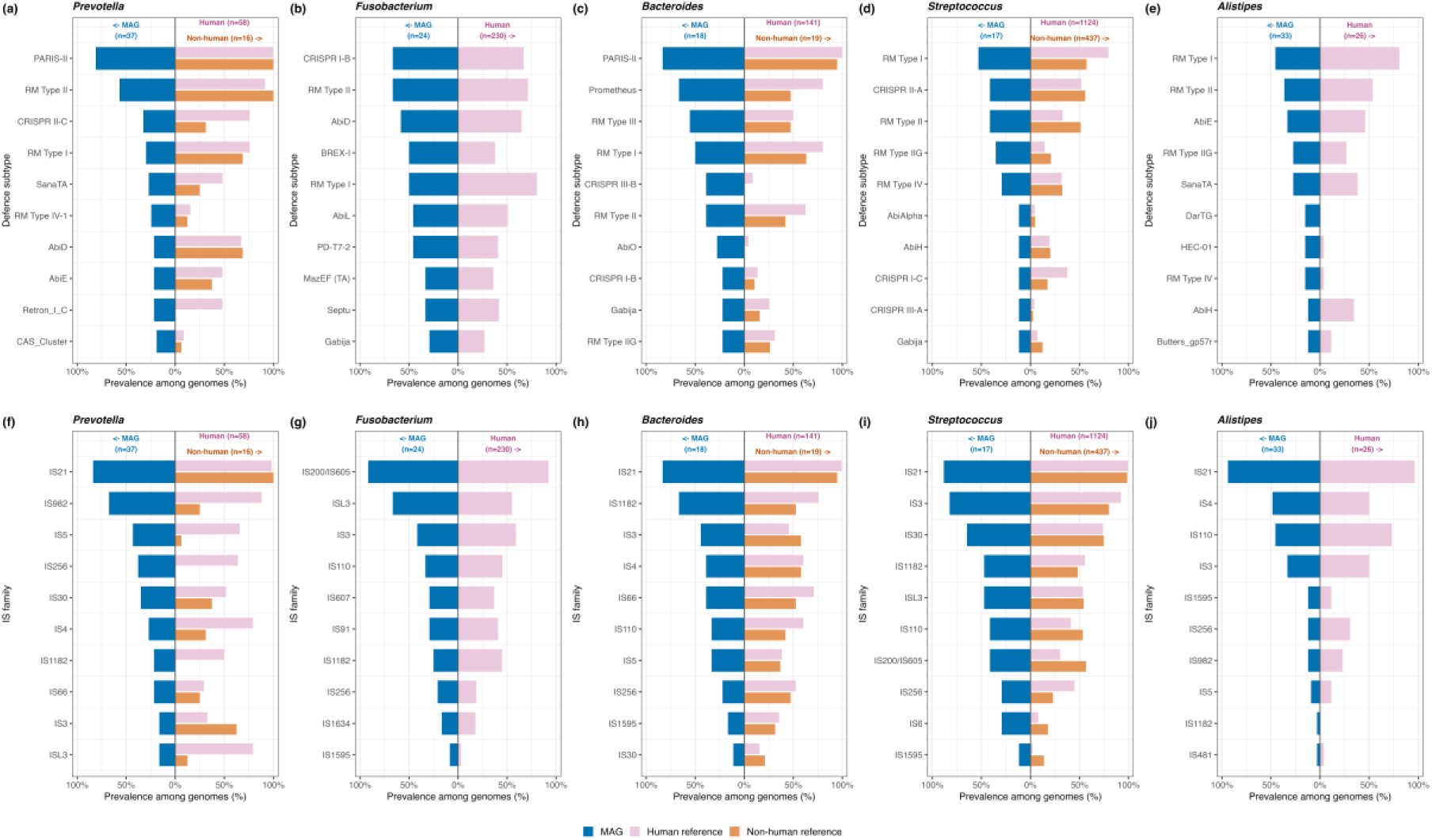
Reference genome validation of MAG based defence system and IS element composition in human-associated genera and the Wirbel *Alistipes* replication cohort. Defence subtype composition is shown in panels a-e, and insertion sequence (IS) family composition in panels f-j. For each genus, the ten most prevalent defence subtypes or IS families among metagenome-assembled genomes (MAGs) are shown. Their prevalence is compared with that in complete NCBI reference genomes of human and, where available, non-human source (chromosomal replicons only). MAG prevalence is shown to the left (blue). Human source (pink) and non-human source (orange) reference-genome prevalence are shown to the right. Panels show four human-associated Hartwig genera (*Prevotella, Fusobacterium, Bacteriodes, and Streptococcus)* and *Alistipes* MAGs from the independent Wirbel colorectal cancer cohort. Only groups with at least 10 genomes are included. *Fusobacterium* and *Alistipes* are compared with human source reference genomes only, because the former had fewer than 10 non-human source reference genomes and the latter had none available. Features were selected according to MAG prevalence only, so defence subtypes or IS families prevalent in reference genomes but absent from the MAG top ten are not displayed.

While screening all 26 defence subtypes shared between the Hartwig human-associated and Wirbel CRC datasets against the reference genome panels pooled across each ecological arm (Table 3), we found that two subtypes, Wadjet-III and AbiQ, recurred across independent cancer-associated datasets while remaining underrepresented in lineage-matched reference genomes. Wadjet-III occurred in 2.3% of Hartwig human-associated MAGs and 6.2% of Wirbel MAGs, but was absent from probable-contaminant MAGs, and undetected in any of the 2,029 reference genomes of the human-associated-arm genera. AbiQ, by contrast, occurred in 5.5% and 6.6% of Hartwig and Wirbel MAGs respectively. It was likewise absent from probable-contaminant MAGs and undetected in the probable-contaminant reference panel. Three further subtypes ranking highly in Table 3, CRISPR II-C, CRISPR III-A, and CRISPR III-D, showed a comparably low probable-contaminant and reference genome prevalence but a less consistent signal across the Hartwig and Wirbel cohorts, leaving their status unresolved. Together, these systems warrant further investigation as candidate markers for distinguishing human-associated from probable-contaminant taxa in future cancer metagenomic pipelines, particularly for genomes with incomplete or unresolved taxonomic classification.

**Table 3.** Candidate defence system markers distinguishing human-associated from probable-contaminant metagenome-assembled genomes (MAGs). Candidates were the 26 defence subtypes shown in Figure 3c, the union of the 20 most prevalent in Hartwig human-associated MAGs (n = 383) and in the Wirbel colorectal cancer (CRC) cohort (n = 785). Of these 11 were absent from Hartwig probable-contaminant MAGs (n = 131), and the ten shown had the lowest maximum prevalence across four reference genome groups (complete RefSeq genomes, chromosome-only tier). Reference genomes are pooled from human-associated arm genera *Prevotella, Fusobacterium, Bacteroides, Streptococcus* and contaminant arm genera *Ralstonia, Bradyrhizobium,* and *Pelomonas*, each split human-source vs. non-human source genomes.

| **System** | **Hartwig human-associated (n=383, %)** | **Hartwig probable-contaminant (n=131, %)** | **Wirbel CRC (n=785, %)** | **Ref. genomes, human-assoc. genera, human-source (n=1553, %)** | **Ref. genomes, human-assoc. genera, non-human (n=476, %)** | **Ref. genomes, contaminant genera, non-human (n=584, %)** | **Ref. genomes, contaminant genera, human-source (n=8, %)** |
|---|---|---|---|---|---|---|---|
| Wadjet-III | 2.35 | 0.00 | 6.24 | 0.00 | 0.00 | 0.51 | 0.00 |
| CRISPR III-D | 7.57 | 0.00 | 1.53 | 2.19 | 0.00 | 0.00 | 0.00 |
| AbiQ | 5.48 | 0.00 | 6.62 | 3.22 | 2.94 | 0.00 | 0.00 |
| CRISPR II-C | 9.66 | 0.00 | 8.15 | 5.54 | 3.15 | 1.54 | 0.00 |
| CRISPR III-A | 7.83 | 0.00 | 6.37 | 6.44 | 2.31 | 0.00 | 0.00 |
| Abi2 | 4.70 | 0.00 | 5.86 | 3.67 | 9.87 | 0.17 | 0.00 |
| CRISPR I-B | 12.79 | 0.00 | 4.20 | 11.14 | 1.68 | 0.00 | 0.00 |
| PARIS-II | 15.14 | 0.00 | 4.71 | 12.81 | 7.14 | 1.20 | 0.00 |
| AbiH | 4.70 | 0.00 | 8.41 | 17.84 | 20.59 | 0.17 | 0.00 |
| RM-Type-III | 15.67 | 0.00 | 13.12 | 17.51 | 27.52 | 10.62 | 12.50 |

Although their biological significance remains to be established, these features represent promising candidate human-associated defence signatures that may contribute to future bioinformatic pipelines leveraging bacteria-phage arms-race dynamics to distinguish biologically significant microorganisms from probable contaminants in cancer metagenomes.

## 8. Discussion

Distinguishing biologically plausible human-associated microorganisms from environmental and reagent-derived contaminants remains one of the central challenges in low-biomass cancer microbiome research(3, 26). In this study, we show that phage-host ecology, defence and anti-defence systems, insertion-sequence (IS) repertoires, plasmids, prophages, and genome architecture may provide complementary ecological signatures that consistently differentiate human-associated MAGs from probable contaminants. Phage-host predictions showed strong concordance with bacterial taxonomy, indicating that associated viral sequences retain ecological information about their hosts, while the recurrence of multiple signatures in the independent Wirbel CRC cohort suggests that these patterns are not specific to a single cancer metagenome dataset.

Hartwig and Wirbel human-associated MAGs exhibited broadly similar defence-system, counter-defence, and IS-family repertoires, whereas probable-contaminant MAGs displayed distinct profiles. However, comparisons with lineage-matched reference genomes indicated that many of the dominant defence systems and IS families were characteristic of the underlying bacterial taxa, suggesting that much of the observed cross-cohort concordance reflects conserved lineage-associated genomic features rather than an ecological signal specific to cancer-derived microbiomes. Nevertheless, a subset of features, including AbiQ and Wadjet-III, recurred across both cancer-associated datasets yet were not prominent within the corresponding reference-genome profiles (Table 3). Wadjet-family systems occur in roughly 6% of sequenced bacterial genomes and act primarily by restricting plasmid-mediated transformation rather than phage infection directly(75). AbiQ is a well characterised abortive-infection system that blocks phage DNA maturation after infection (76–78). Their relevance to cancer-associated microbial ecology therefore remains to be established through lineage-matched comparisons and experimental validation. Once a cancer-associated bacterial target has been independently validated, its defence profile could be incorporated into phage-selection strategies alongside receptor availability, host range, and experimentally determined infectivity.

Plasmid and prophage analyses provided additional, although context-dependent, ecological information. Human-associated MAGs from both cohorts displayed similarly low plasmid-associated sequence burdens, whereas probable-contaminant MAGs were strongly enriched in plasmid-associated sequences. In contrast, prophage-associated features were less consistent between cohorts. Hartwig human-associated MAGs contained a higher proportion of temperate-associated and gene-rich proviruses than Wirbel CRC MAGs, suggesting that prophage architecture may be influenced more strongly by tissue-associated versus luminal microbial ecology than by cancer association alone. Together, these findings indicate that different components of the mobilome provide complementary ecological information, with some signatures showing broad reproducibility across cancer-associated datasets and others appearing more dependent on habitat-specific ecological context.

### Implications for defence-informed phage therapy

Phages are increasingly being explored as precision tools for targeting cancer-associated bacteria while minimising disruption of the wider microbiome(22). This is particularly relevant in oncology, where broad-spectrum antibiotics can perturb commensal communities and potentially influence treatment responses(79–82). Preclinical studies targeting bacteria such as *Fusobacterium nucleatum* illustrate the potential of phage-based approaches to reduce tumour-promoting microbial effects(83–86).

Our reference-genome analysis indicates that many defence systems detected in cancer metagenomes are characteristic of the dominant bacterial lineages, including *Fusobacterium*. However, an important limitation is that the available reference genomes for several retained genera were predominantly human associated; suitable complete non-human references were limited or unavailable (*Fusobacterium* (4/239), *Leptotrichia* (0/22), and *Alistipes* (0/27). The observed similarity between human-associated MAGs and human-associated reference genomes may therefore reflect both phylogenetic relatedness and shared host ecology. Comparisons with closely related strains from contrasting ecological sources will be needed to distinguish lineage-specific from host-or cancer-associated defence signatures.

The high prevalence of temperate-associated prophage signatures in some Hartwig MAGs also has implications for therapeutic design. Phages intended for bacterial depletion should ideally be strictly lytic and screened for lysogeny, virulence, antimicrobial-resistance, and horizontal-gene-transfer determinants. Integration of high-throughput phage-host screening, functional genomics, and predictive modelling could therefore help combine bacterial defence architecture with phage host range and safety characteristics to guide more rational phage selection.

### Limitations and future directions

We acknowledge several limitations that should be considered when interpreting these findings. MAG reconstruction introduces inherent uncertainty, particularly in low-biomass tumour metagenomes where assemblies are often fragmented. This can reduce the recovery of complete defence islands, prophages, plasmids, and repetitive IS elements. Large plasmids may also be split across multiple contigs, while repetitive IS elements can collapse during assembly. For this reason, absolute estimates of MGE abundance should be interpreted with caution.

Another challenge was no negative-control sequencing data were available from the original studies, preventing direct comparison with conventional contamination-detection approaches. The Hartwig MAGs were analysed as deposited by Battaglia et al. without access to the underlying sequencing reads. The Wirbel MAGs were assembled in this study from public stool metagenomes for which no reagent or extraction blanks were available. Battaglia et al.(2) assessed read-level contamination in the Hartwig cohort using matched blood-derived whole genome sequencing as a negative control, but control-based decontamination tools such as decontam(87), which compare taxon prevalence between samples and blanks, could not be applied or benchmarked here. Our screen therefore relied on taxonomic assignment, host ecology prediction, and reference genome source metadata, which are independent of sequencing controls but cannot fully exclude reagent derived lineages that also occur as genuine human commensals.

A related limitation concerns the defence detection tools themselves. DefenseFinder relies on homology-based detection against predefined system models and are consequently prone to missing highly divergent or entirely novel defence systems that lack recognisable similarity to characterised examples. Given the unusual selective pressures bacteria may encounter within tumour-associated microbial communities, it is plausible that additional yet to be characterised defence and counter-defence systems remain undetected in these datasets. Future approaches based on machine learning, protein language models, or other sequence-independent methods may help identify such systems. Recently developed tools of this kind have already discovered, and experimentally validated dozens of previously unrecognised defence mechanisms missed by homology search alone (88–90), making this a promising direction if tumour-associated microbiomes do harbour defence systems beyond those captured by current pipelines. Furthermore, several classes of defence-associated elements, including non-coding RNAs, transfer RNAs (tRNAs), and small signalling molecules, have increasingly been implicated in phage defence (91) are not comprehensively captured by current detection pipeline. Future studies should therefore extend beyond protein-coding defence genes to investigate the potential contribution in cancer metagenomes. In addition, comprehensive characterisation of CRISPR arrays and spacer content may provide further insights into historical phage exposure, host-virus interactions, and the evolutionary dynamics of human-associated microbial communities.

A further limitation is the difference in biological source between the two cancer cohorts. Hartwig MAGs were reconstructed from metastatic tissue biopsies, whereas Wirbel MAGs were derived from CRC stool. Although the Wirbel cohort provides valuable external validation, the two datasets represent distinct microbial habitats and are therefore not directly comparable for all phage-related features. In particular, the differences observed in prophage lifestyle may reflect the contrasting ecology of tissue-associated and luminal communities rather than cancer status itself. Future studies incorporating matched tumour tissue, adjacent non-tumour tissue, stool, blood, and appropriate environmental controls would provide a stronger approach for separating cancer-specific microbiome from anatomical niche effects.

Larger, carefully matched datasets will be required to determine whether defence–mobilome features are genuinely associated with cancer rather than with bacterial lineage or host ecology. Our reference-genome comparisons showed that many defence systems and IS-family patterns detected in cancer-derived MAGs were also characteristic of the corresponding human-associated lineages. This study also lacked a non-cancer comparator. Future studies should therefore compare cancer and non-cancer microbiomes within species-or strain-matched groups and, where possible, from the same anatomical site. Controlled-access datasets available through dbGaP could support such analyses across multiple tumour types using a standardised workflow for host depletion, assembly, MAG reconstruction, defence profiling, and MGE characterisation. Integration with tumour type, anatomical site, disease stage, treatment history, immune features, sequencing centre, and clinical outcome would help distinguish lineage- and niche-associated variation from defence–mobilome features that may be specifically associated with the tumour environment. Ultimately, sufficiently large and well-controlled datasets could support predictive models for matching phages to validated tumour-associated bacterial targets, providing a more rigorous basis for defence-informed and potentially personalised phage therapy.

## Conclusion

In this study, we demonstrate that integrating phage–host ecology such as defence & anti-defence profiles, insertion sequences, plasmids, prophages, and genome architecture provides a multilayered strategy for assessing the ecological plausibility of metagenome-assembled genomes recovered from complex host-associated datasets. Human-associated MAGs from the Hartwig cohort showed broad concordance with those recovered from the independent Wirbel colorectal cancer dataset, whereas probable contaminants exhibited distinct defence and mobilome profiles. Although many of the observed signatures largely reflected underlying bacterial lineages, a subset of features, including AbiQ and Wadjet-III, recurred across independent datasets while remaining rare in reference genome collections and absent from probable contaminants.

Rather than serving as direct evidence of cancer-specific microbial adaptation, these findings highlight the value of defence and mobilome signatures as complementary ecological markers for evaluating MAG authenticity and distinguishing probable contaminants from biologically plausible host-associated genomes. Our results further underscore the need for larger, geographically diverse, and experimentally validated datasets to disentangle lineage-associated patterns from potential disease-associated signals.

More broadly, this work illustrates how phage–host interaction features can complement conventional quality-control metrics in metagenomic studies. As reference databases expand and experimental validation becomes increasingly feasible, ecology-informed frameworks such as this may improve confidence in MAG interpretation and support more robust investigations of microbial communities across health and disease.

## 9. Author statements

### 9.1 Author contributions

V.M. contributed to conceptualization, formal analysis, methodology validation, visualization, writing of the original draft, and review and editing of the manuscript. S.S. contributed to conceptualization, data curation, formal analysis, investigation, methodology, resources, supervision, writing of the original draft, and review and editing of the manuscript. V.R. contributed to formal analysis, writing of the original draft, and review and editing of the manuscript.

### 9.4 Ethical approval

Not applicable

## 9.5 Acknowledgements

Not applicable

