## Supplemental Methods and Figures for "Artefact or ecology? Integrating bacteria-phage defence and mobilome signatures to distinguish human-associated MAGs from probable contaminants in low biomass cancer metagenomes"

**Contents:** Supplementary Methods; Supplementary References; Figure S1; Figure S2.

**Supplementary Methods: Processing of the Wirbel colorectal cancer cohort-level**

Paired end reads of all 60 colorectal cancer samples of the German cohort of Wirbel et al. [1] were downloaded from ENA (PRJEB27928, 22 samples; PRJEB6070, 38 samples [2]). Human reads were removed with Hostile (v2.0.2) [3] against a phage-masked T2T-CHM13 reference [4] with HLA sequences, followed by Kraken2 [5] classification against the PlusPF-16 database to remove residual human reads. Reads were quality filtered with fastp (v0.23.4; minimum base quality Phred 20, minimum length 50 bp, low complexity filtering at threshold 30, and deduplication) [6].

Each sample was assembled with MEGAHIT (v1.2.9; k-mers 21-141 in steps of 12, minimum contig length 500 bp) [7]. Reads were mapped back to contigs with Bowtie2 [8] to estimate coverage, and contigs were binned with MetaBAT2 [9]. Of the 4,150 resulting bins, 4103 were assessed after excluding MetaBAT2’s unbinned, too-short and low-depth bins. Completeness and contamination were estimated with CheckM2 (v1.0.1, database v2) [10], and the 785 bins with ≥90% completeness and ≤5% contamination were retained as high-quality MAGs. These came from 55 samples; the remaining 5 yielded on high-quality bin because of high contamination or low sequencing depth. Taxonomy was assigned with GTDB species representative (ani_rep) which resolved 783 of the 785 MAGs.

MAGs were annotated with Bakta (v1.12.0, database v6; --meta mode) [13]. Defence and anti-defence systems were identified in a single run of DefenseFinder (v2.0.1 models v2.0.2) [14,15], viral and plasmid sequences with geNomad (v1.11.2) [16], viral quality with CheckV (database v1.5) [17], phage genes with Pharokka 9v1.9.1) [18], IS elements with ISEScan 9v1.7.3) [19] and phage hosts with iPHoP (database Jun_2025_pub_rw) [20]. MAGs for which ISEScan or iPHoP produced no output after re-running were recorded as carrying no IS elements or host predictions respectively. MAGs were classified with the same three step workflow as the Hartwig cohort (taxonomic screen, iPHoP host profile and literature review of flagged lineages; Table 2).

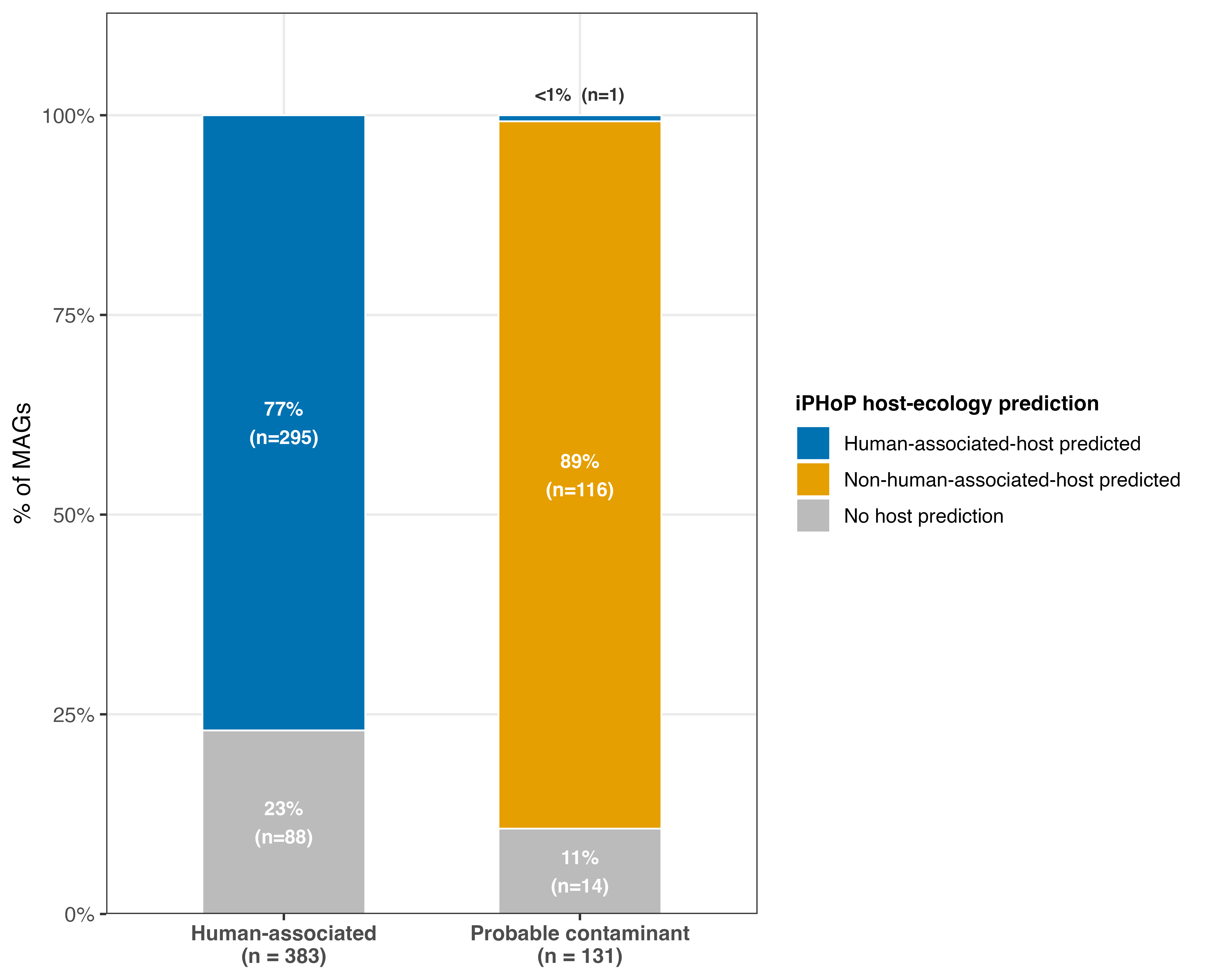

**Figure S1: Predicted phage host ecology: Hartwig human-associated vs probable-contaminant MAGs.**

Each metagenome-assembled genome (MAG) is classified by the ecology of its predicted phage hosts. iPHoP host predictions were classified as human-associated or not by host phylum and genus (Table S3). Each MAG is then assigned one of three categories: human-associated-host predicted (at least one human-associated host prediction), non-human-associated-host predicted, or no host prediction. Human-associated MAGs (n = 383) carry a human-associated-host predicted phage in 77% of cases (295/383), versus under 1% (1/131) for probable-contaminant MAGs (n = 131) (Fisher’s exact test, odds ratio (OR) = 430, p = 5.7×10⁻⁶¹).

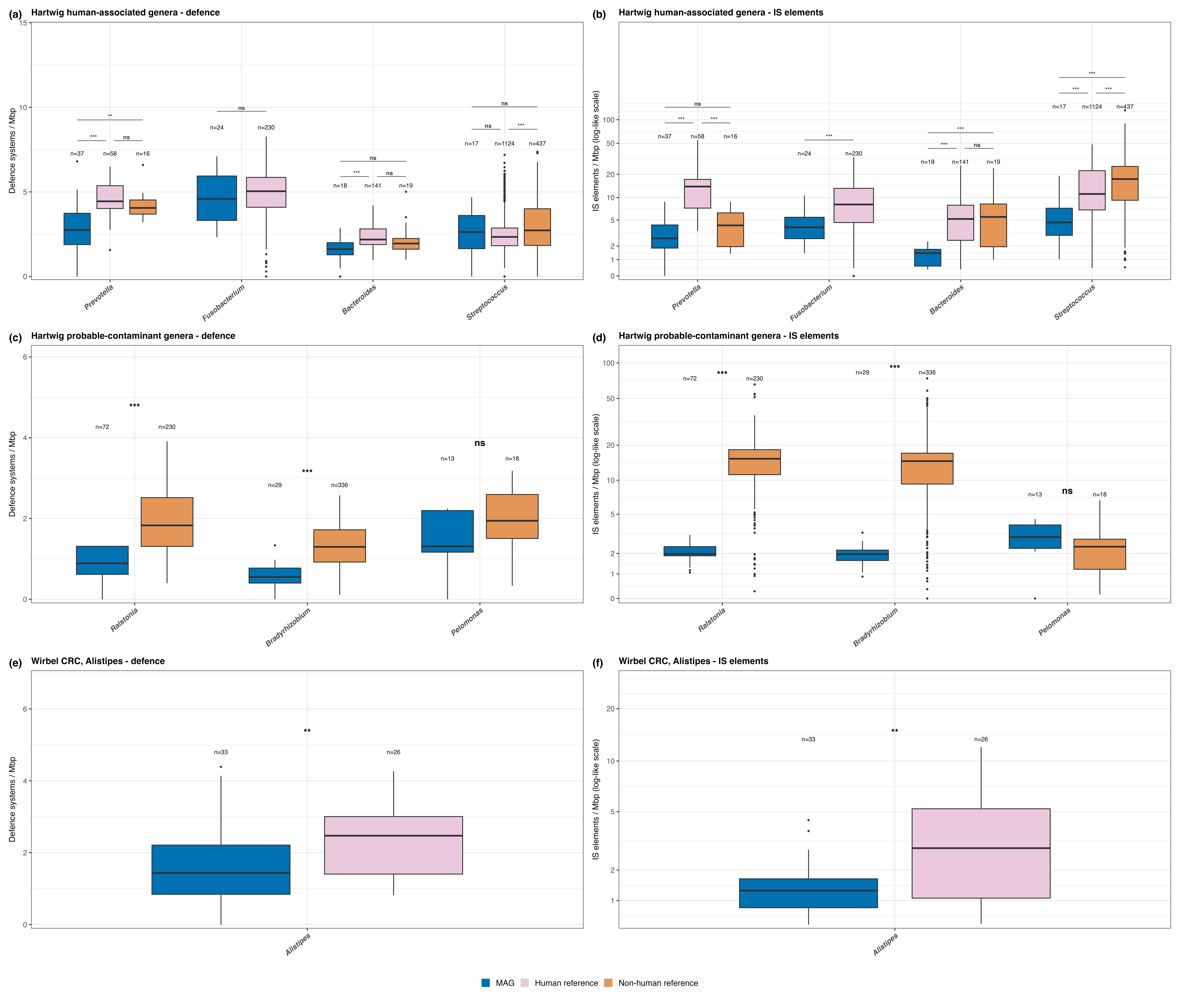

**Figure S2: Reference-genome validation of MAG based defence system and IS element burden per Mbp of genome.**

Defence system burden (A, C, E) and insertion sequence (IS) element burden (B, D, F) are shown for metagenomes-assembled genomes (MAGs) and complete NCBI reference genomes from the same genus. For reference genomes, defence systems and IS elements were counted on chromosomal replicons only and normalised per Mbp of chromosome. IS burden in shown on a pseudo-log scale. Boxplots show the median, interquartile range (IQR) with 1.5 x IQR whiskers, and n is given for each group. Asterisks indicate two-sided Mann-Whitney U tests with Benjamini-Hochberg correction across genera within each comparison (*p < 0.05, **p < 0.01, ***p < 0.001; ns, not significant). Only groups with at least 10 genomes are shown. (A, B) Human-associated Hartwig genera (*Prevotella*, *Fusobacterium*, *Bacteroides*, and *Streptococcus),* compared with human source and non-human source reference genomes. *Fusobacterium* is compared with human source reference genomes only, as fewer than 10 non-humn source genomes were available. (C, D) Probable-contaminant genera (*Ralstonia*, *Bradyrhizobium*, and *Pelomonas)* compared with non-human source reference genomes. (E, F) Wirbel colorectal cancer (CRC) *Alistipes* MAGs compared with human source reference genomes.
